# Near-atomic structure of the PorKN rings, disulfide bonded to PorG and bound to Attachment Complexes, provide mechanistic insights into the type IX secretion system

**DOI:** 10.1101/2025.03.18.644022

**Authors:** DG Gorasia, E Hanssen, M Mudaliyar, CJ Morton, S Valimehr, C Seers, L Zhang, D Ghosal, PD Veith, EC Reynolds

## Abstract

The Type IX Secretion System exports proteins across the outer membrane (OM) of bacteria in the Bacteroidota phylum, however, the mechanistic details remain unknown. Here, we present a ∼3.5Å cryo-EM structure of the periplasmic rings comprising 32-33 subunits each of PorK and PorN. Additionally, we show the presence of a critical disulfide bond between PorK and the PorG OM protein that is essential for protein secretion and demonstrate that the Attachment Complexes bind to and are localized above the PorKN rings. Overall, each ring resembles a cogwheel with PorN forming cog-like projections on the periplasmic side and the flat surface of PorK orienting towards the OM. Given these results, we propose that the PorLM motor drives the rotation of the PorKN cogwheel together with PorG and associated Attachment Complexes, potentially providing the energy to complete protein secretion and the coordinated cell surface attachment of the secreted cargo.

## Introduction

Periodontitis is a prevalent, chronic inflammatory disease associated with a dysbiotic subgingival plaque that results in the destruction of the tooth’s supporting tissues and is additionally linked with a range of systemic diseases like diabetes, cardiovascular diseases, certain cancers and dementia [1, 2]. *Porphyromonas gingivalis,* an obligate anaerobe, is strongly associated with severe periodontitis at sites refractory to treatment and is now recognized as a keystone pathobiont of this disease [3–5]. *P. gingivalis* uses the Type IX Secretion System (T9SS) to translocate at least 35 cargo proteins to the cell surface, including hallmark virulence factors such as the gingipain proteases RgpA, RgpB, and Kgp [6, 7]. The T9SS secreted cargo proteins play an important role in virulence of the bacterium, including biofilm formation, nutrition acquisition, antibiotic resistance, adhesion and degradation of host proteins [8, 9].

The T9SS is exclusively present in members of the *Bacteroidota* phylum [10] [11]. The T9SS cargo proteins have an N-terminal signal peptide that facilitates export across the inner membrane by the Sec system and have a conserved C-terminal domain, referred to as the CTD signal, that enables them to pass through the outer membrane (OM) via the T9SS [7, 12–14]. Once on the surface, CTD signals are removed and replaced with anionic-lipopolysaccharide (A-LPS), thus anchoring the proteins to the cell surface to form an electron dense surface layer (EDSL).

Besides regulatory proteins, the T9SS is composed of at least 18 proteins in *P. gingivalis*, namely: PorK, PorL, PorM, PorN, Sov, PorT, PorU, PorW, PorP, PorV, PorQ, PorZ, PorE, PorF, PorG, Plug, PorD, PorA [6, 8, 15–17]. Recent studies have begun to shed light on the structure and function of some of these components (**Fig 1A**). The PorL-PorM motor complex is anchored to the inner membrane and powered by the proton motive force [18, 19{Hennell James, 2021 #20]}. The PorM rotor extends into the periplasm to drive both secretion and motility[20]. Additionally, *in-situ* structural studies of the T9SS and bacterial two-hybrid studies have shown an interaction between PorM and PorN [18, 21]. PorK and PorN form large ring-shaped structures in the periplasm that are tethered to the OM by the lipidation of the PorK N-terminus and by association with an 8-stranded OM β-barrel protein, PorG[19, 21]. Evidence from cross-linking studies indicated that a periplasmic loop of PorG is in close proximity to both PorK and PorN [19]. SprA (an orthologue of *P. gingivalis* Sov) has been shown to form a large 36-stranded β-barrel that is either bound to PorV or Plug and is the OM translocon through which the T9SS cargo proteins are secreted [22]. We demonstrated that the Sov interactome in *P. gingivalis* included PorV, PorA, Plug, PorW, PorD, and the PorKN rings, proposing that PorW forms a bridge between the PorKN rings and the Sov translocon.[23]. Recent findings in *F. johnsoniae* validated our results, where an extended-translocon including SprE (PorW orthologue), PorD, and trapped substrates was purified from the *gldL* (*porL*) mutant. The accumulation of substrates in the translocon when energetic input was removed suggested that energy is required to release PorV-bound substrates from the translocon [24]. PorU and PorZ are anchored to the OM via binding to the PorV and PorQ OM β-barrel proteins respectively, and together the four proteins form the Attachment Complex [25]. PorU is a sortase that cleaves the CTD signal on the cell surface and swaps it with anionic-lipopolysaccharide (A-LPS) provided by PorZ, thereby anchoring cargo proteins onto the cell surface [26–28].

**Figure 1:**
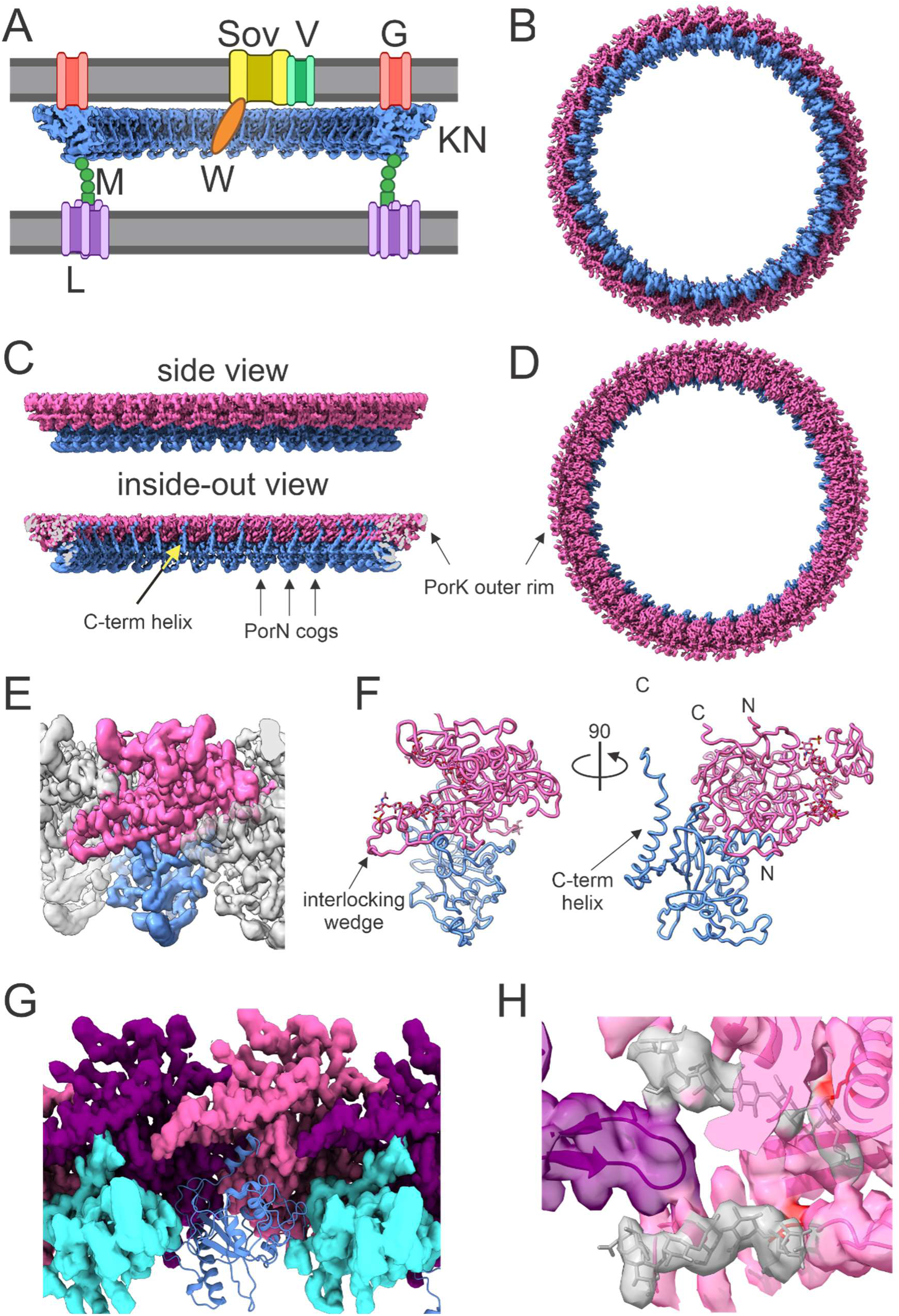
Structure of the PorKN ring. (A) Cartoon representation of the T9SS architecture showing the location of PorKN rings and PorG, PorLM motor, and the Sov translocon bound to PorV and PorW. (B) A bottom view of 33-fold symmetry cryo-EM volume of the PorKN ring (Pink: PorK, Blue: PorN). (C) Side views of the PorKN ring showing the flat upper surface and an outer rim formed by PorK with cog-like projections formed by PorN. (D) Top-view of the PorKN ring. (E) Enlarged views of the outer surface of the PorKN ring showing the fitting of PorK (pink) and PorN (blue) and interlock of PorK subunits. (F) Ribbon fold structures of PorK (pink) and PorN (blue), front and side views showing the glycans on PorK. (G) Inner ring cross section showing the interface of PorN with the cavity formed by two PorK subunits. PorK (purple and pink) surface rendered. PorN (blue) in surface rendered and ribbon representation. (H) Detailed structure of the two glycans on PorK.

Although, the PorKN ring and its association with PorG was identified nearly a decade ago, their precise role remains unclear. The proteins that comprise the Attachment Complex have been identified, but the exact location of this complex in relation to the other T9SS components is unknown. Here, we present a high-resolution structure of the PorKN ring, offering new insights into their function. We also demonstrate that PorG is associated with PorK through a disulfide bond, which is crucial for the function of the T9SS and show that the Attachment Complexes are associated with, and co-localised *in-vivo* above the PorKN rings providing insight into the co-ordinated mechanism of T9SS secretion and attachment in *P. gingivalis*.

## Methods

### *P. gingivalis* culture conditions

*P. gingivalis* strains were grown on solid medium (TSBHI agar) containing trypticase soy agar (40 g/L), brain heart infusion (5 g/L), 5% (v/v) lysed defibrinated horse blood, cysteine hydrochloride (0.5 g/L), and menadione (5 mg/mL) or in TSBHI broth (25 g/L tryptic soy, 30 g/L brain heart infusion, 0.5 g/L cysteine, 5 mg/L haemin and 1 mg/L menadione) under anaerobic conditions as described previously [29, 30]. The appropriate antibiotic was included in the media to select for the relevant antibiotic marker of the strain but omitted in the final harvested liquid culture.

### Construction of *porG* deletion mutant

The *porG* deletion mutant in the W50 strain was generated by double cross-over recombination using the suicide plasmid pΔPorG-ermF that contained upstream and downstream regions of *porG* (PG0189) directionally cloned adjacent to an erythromycin resistance cassette (*ermF*). The upstream and downstream regions of *porG* were amplified by PCR using W50 genomic DNA as a template and the following primer pairs: Upstream-F (ACGC<u>GCATGC</u>CCACAGAACAAGCCGAAGGTATAA (SphI site underlined)/ Upstream-R (ACGC<u>CCATGG</u>TGAGCAGCTATCCGGCCGTTAC) (NcoI site underlined) and Downstream-F (ACGC<u>CTGCAG</u>GGATGCGCTGACGGATAAAAGTG) (PstI site underlined)/ Downstream-R (ACGC<u>ATGCAT</u>ACCATCCATGACCAAAGCGATATG) (NsiI site underlined). The PCR amplicons were digested with the appropriate restriction endonucleases and sequentially cloned upstream and downstream of *ermF* in pAL30 [31]. Once the sequence integrity of the cloned DNA was confirmed pΔPorG-ermF was linearised using NsiI and electroporated into *P*. *gingivalis* W50 with transformants selected on TSBHI agar plates supplemented with 10 mg/L of erythromycin. The correct homologous recombination of transformants was verified using PCR with primer pair TACTATTATGCCACAGAAC and GACAACCACCCGACTTTGAA. The absence of PorG protein was verified by mass spectrometry.

### Construction of PorG complementation strain

To construct the *porG* complementation plasmid, pKD713-PstI that contains a recombinant cassette of the *fimA* neighbouring gene regions and the *tetQ* gene selection marker (a gift from Dr Mikio Shoji) was utilized. DNA spanning from a 3′ region of the PG0188 locus to *porG* was PCR-amplified using primers -ACGC<u>CTGCAG</u>CCATGGTTTGGCCACAGGGAA and - ACGC<u>CTGCAG</u>GCACTATAACTGCTGACTGGAG (PstI site underlined). The amplicon was cloned into the pKD713-PstI through PstI digestion and ligation to create pKD713-PorG. Once it was verified that the cloned DNA sequence was correct, pKD713-PorG was linearised using NotI and introduced into the *porG* mutant cells by electroporation and plated on TSBHI agar supplemented with 1 mg/L tetracycline to select complementation clones. Insertion of *porG* into the *fimA* locus was verified by genomic PCR using primers corresponding to *porG*, the tetracycline resistance gene (*tetQ*) and the *pg2133* locus.

### Construction of targeted PorG^Cys232-Ser^ *P. gingivalis* mutant strain

A series of PCR and SOE-PCR reactions were conducted to produce *pg0189* Cys232Ser codon mutation and non-mutated control recombination cassettes. The *pg0188* open reading frame 3’ overlaps the 5’ of the *pg0189* open reading frame by 14 nucleotides so the recombination marker *ermF* was placed 3’ to *pg0189* to not disrupt the putative operon. Template DNAs were pAL30 [31] (PCR-1) or *P. gingivalis* W50 genomic DNA (PCR-2, PCR-3 and PCR-4). PCR-1 used primers ermFprom_for (GATAGCTTCCGCTATTGCTT) and ermF-Rev (GACAACCACCCGACTTTGAA) to produce *ermF* with the *ermF* promoter. The PCR-2 amplicon used a forward primer with a fusion of ermF 5’ and *pg0189-pg0190* intergenic nucleotides. The PCR-2 primers were ermF_pg0190_For, <u>AAGCAATAGCGGAAGCTATC</u>CCTCGAAACATTCTAAGGGAAT (*ermF* nucleotides underlined) and a reverse primer that annealed within *pg0190,* Reverse_3_PstI, ACGC<u>CTGCAG</u>CTTCCGGACGATTCCAATTCT (PstI site underlined). PCR-3 and PCR-4 used forward primer that annealed to *pg0189*, Forward Oli_1_SphI (ACGC<u>GCATGC</u>GGCCGGGTTGTCTTTGGGTT, SphI site underlined) and respective reverse primers that annealed to *pg0189* and had an *ermF* tag at 5’ (underlined below), Reverse-1A-WT (<u>TTCAAAGTCGGGTGGTTGT</u>CCCCTATTGTTTATTACAAAAAGTC) and Reverse-1B-Ser (<u>TTCAAAGTCGGGTGGTTGT</u>CCCCTATTGTTTATTACTAAAAGTC). PCR-1 and PCR-2 were denatured and annealed by the homologous nucleotides, extended, then amplified with external primers ermF-Rev and Reverse_3_PstI to produce SOE-1 product. SOE-1 and either PCR-3 or PCR-4 were denatured and annealed, extended, then amplified with external primers Forward Oli_1_SphI and Reverse_3_PstI to produce SOE-2 and SOE-3 respectively. SOE-2 and SOE-3 were digested using SphI and PstI and ligated to pAL30 vector. After verification of the fidelity of the insert DNA by sequencing the plasmid was digested with SacI and transformed into *P. gingivalis* W50 by electroporation. Erythromycin-resistant transformants were selected from TSBHI agar plates and correct integration into the genome was determined using PCR with primer pair Forward Check (GCGGATTTTAAGGTCGGCTT) and Reverse-Check (AGCATCTCATTCATGATAGC).

### Gingipain activity assay

Arg- and Lys-specific proteolytic activity was determined using the synthetic chromogenic substrates N-benzoyl-DL-arginine-4-nitroanilide hydrochloride (BapNA) and N-(*p*-tosyl)-Gly-Pro-Lys 4-nitroanilide acetate salt (GPKNA) (Sigma Aldrich), respectively. Briefly, *P. gingivalis* in broth culture were pelleted at 8000g for 10 min then resuspended in a buffer containing 50 mM Tris-HCl, pH 7.4, 5 mM CaCl_2_ and 150 mM NaCl. The supernatant (culture fluid, CF) containing vesicles were also collected. Assays were conducted in 96 well microtiter plates with 5.6 x10^6^ cells or 140 µL of CF per well to which a final concentration of 20 mM L-cysteine, pH 8.0 was added. After the mixture was incubated at 37°C for 10 min, a final concentration of 1 mM substrate (BapNA or GPKNA) was added to it and substrate cleavage was determined by measuring the absorbance at 405 nm at 10 s intervals for ∼ 60 min at 37°C using a PerkinElmer 1420 Multilabel Counter VICTOR3™. For culture fluid (CF) activity assay, 140 µL of each sample was used instead of the cells.

### SDS-PAGE and western blots

Protein samples were mixed with SDS-loading dye and analysed on NuPAGE Bis-Tris precast gels electrophoresed using MOPS SDS running buffer. The gels were stained overnight with 0.1% w/v Coomassie Blue G-250 in 17% w/v ammonium sulphate, 34% v/v methanol and 3% v/v o-phosphoric acid. For western blotting, protein samples were resolved on SDS-PAGE gels and transferred onto a nitrocellulose membrane. The membrane was blocked with 5% skim milk powder in PBS-tween (0.05% (v/v) Tween-20 in PBS). The membranes were probed with mAb-1B5 (a kind gift from Professor M.A Curtis), PorK or PorN primary antibodies followed by goat anti-mouse or anti-rabbit HRP conjugated secondary antibodies. The signal was developed using SuperSignal West Pico chemiluminescent substrate and visualized with a LAS-3000 imaging system.

### Cell envelope (CE) preparation

The cell envelope was prepared as described in Glew et al [29]. Briefly, *P. gingivalis* cells were washed and resuspended in 0.5x PBS, 5 mM MgCl_2_ in a 50 mL tube and placed in a beaker containing an ice/water mixture and sonicated using an ultrasonic processor (model CPX 750, Cole Parmer) fitted with a 6.5 mm tapered microtip. The amplitude was set to 40%, pulser to 1 s on, 2 s off for a total of 30 min. The CE was collected by centrifugation at 48400 *g* for 20 min and washed in 1x PBS before storing in -80°C for further analysis.

### Blue native PAGE immunoblot and co-immunoprecipitation

Blue native PAGE immunoblot and co-immunoprecipitation were conducted using cell lysates in buffers containing 1% DDM as described by Gorasia et al, [23, 32, 33].

### Purification of PorKNG complex

PorKNG complexes were purified as per our published protocol [19]. For the complex preparation under reducing conditions, 5 mM dithiothreitol (DTT) was added to the cell lysate and 1 mM DTT in all the subsequent steps. For the cryo-EM single particle analysis, 1 M urea was added to the purified sample immediately before application to the EM grid.

For the crude preparation, the *P*. *gingivalis* cells (ABK-(*P. gingivalis* cells lacking gingipains) and *porG* mutant) were lysed and digested with lysozyme as per our published protocol [19] and the pelleted complexes were resuspended in 50 mM Tris-HCL, pH 7.5, 500 mM NaCl, 1% DDM and 5 mM EDTA. Following centrifugation at 17,000 g for 10 min, the resulting supernatant was layered on a 30% sucrose cushion and centrifuged at 35,000 rpm for 2 hours at 4°C in a TLS-55 rotor (Beckman Coulter). The pellet was resuspended in 10 mM Tris-HCl, pH 7.5, 50 mM NaCl and 0.5% DDM and stored at 4°C.

### Electron microscopy

For negative staining of protein complexes samples were adsorbed onto glow discharged Formvar-carbon films supported on 400 mesh copper grids. Samples were negatively stained with aqueous uranyl acetate and viewed under an FEI Tecnai F30 operated at 200kV equipped with a CETA 4kx4k camera (FEI, NL). Micrographs were obtained under low-dose conditions (∼20 e^−^/Å^2^) with defocus values of ∼ 2 μm. For cryo-electron microscopy of protein complexes and *P*. *gingivalis* cells, samples were applied onto a glow discharged Quantifoil (respectively R1.2/1.3 and R3.5/1) holey carbon film mounted on a 200 mesh copper grid. The grid was blotted for 4 seconds with a blot force of -1 using a Vitrobot Mark IV (Thermo Fisher Scientific) set to 22°C and 100% humidity and plunge frozen into liquid ethane. For cells, transmission electron microscopy was carried out under cryogenic conditions using a FEI Tecnai F30 operated as described above. Micrographs were obtained under low-dose conditions (∼25 e^−^/Å^2^) with defocus values of ∼2 μm. For protein complex, 1 M urea was added to the complex to limit strong self-aggregation and cryo-EM data collection was carried out on a Titan Krios G4 cryo-electron microscope equipped with a K3 detector and a BioQuantum imaging filter (Gatan) at 300 kV in counting mode. Movie stacks were collected at a nominal magnification of 64,000 ×, a pixel size of 1.32 Å and a defocus of -0.6 to -2 μm with a fluency of 13.5 e^−^/pixel/s. Each movie was a result of 5.34 s exposures with a total accumulated electron exposure of 54 e^−^/Å^2^ which were fractioned into 40 frames. Automated data collection in AFIS mode was performed using EPU (Thermo Fisher Scientific). An energy filter slit width of 10 eV and a 50 μm C2 condenser aperture were used during imaging.

### Cryo-EM Data Processing

Movies were collected (8,909), motion-corrected and CTF fitted using CryoSPARC Patch motion correction and Patch CTF jobs [34]. Manual picking was used to pick ∼200 particles, subject them to class average with 3 classes corresponding to top, side and tilted views. Picking using these templates yielded 556,522 initial particles from which 121,937 final particles were selected using two rounds of 2D classification. Ab initio modelling with 50,000 particles was carried out and output 4 model. The top two models were used in heterogenous refinement with the 121,937 particles. 77,661 particles were selected and subjected to homogenous refinement in C1 then C2. Symmetry expansion and subtraction on one of the ring was carried out. The resulting particles were subjected to C1 homogenous refinement and 3D classification in C1, 47.3% resulted in a map with 33 repeats and 52.3 % in a ring with 32 repeats. Both maps were subjected to C1 refinement then respectively C33 and C32 refinement. Particles were further cleaned up with a round of 2D classes and a final round of local refinement (masking the detergent) was carried out in respectively C33 (3.57 Å, EMD-48741) and C32 (3.63 Å, EMD-48722).

### Structure refinement

Initial model of PorK and PorN were generated using Alpha Fold3 with three copies as a hexamer (three copies of the PorK-N dimer) and fitted to the experimental map with manual modification in Coot [35] and real space refinement with Phenix [36] with statistics reported for the central dimer of the modelled hexamer (**Supplementary Table 1**). Complete PorKN ring models were generated by duplication of the refined dimer structure using C33 symmetry (PDB: 9MYJ)

### Immunogold labelling and cryo-electron tomography

PorZ antibodies were generated by Genscript against amino acid residues 29-776 of *P. gingivalis* PorZ. *P. gingivalis* cells were labelled with PorZ antibodies as detailed in Chen et al. [30] with the following modifications. PorZ antibodies were used at a concentration of 30 µg/mL and three washes were included after the incubation with anti-rabbit secondary gold (6 nm) antibody. The immunogold treated positive and appropriate negative control samples were vitrified using a Vitrobot Mark IV (Thermo Fisher Scientific). 4 μL of the sample was pipetted on a glow discharged Quantifoil extra thick copper EM-grids (R2/2, 200 mesh, Electron Microscopy Sciences). The excess liquid was blotted (blot force 7, blot time 7 seconds) using a Whatman filter paper and plunge frozen into liquid ethane.

Data was collected on Titan Krios G4, 300 keV FEG cryo transmission electron microscope, equipped with a Gatan K3 direct electron detector equipped with an energy filter set to 20 eV. The tilt series was acquired in movie mode from –60° to +60° at 3° intervals with a total electron dose of 120 e^-^/ Å^2^, defocus of –8 μm and a pixel size of 3.4 Å. Subsequently, tilt series were aligned using IMOD (Version 4.11.5) [37]. The 2K binned micrographs were reconstructed into 500 nm thick 1K SIRT tomograms by Tomo3D (Version 2.2) [38].

### In gel digestion and mass spectrometry

In-gel digestions and LC-MS/MS analysis were performed as essentially described in [19, 23, 27]. Relative abundances of proteins were quantified by MaxQuant (version 1.5.3.30) [39]. Raw MS/MS files were searched against the *P. gingivalis* W83 or ATCC 33277 database. The default MaxQuant parameters were used, except the label-free Quantitation (LFQ) minimum ratio count was set to 1 and the match between runs was selected. MaxQuant normalized the data set as part of data processing.

To identify the disulfide bond, raw MS/MS files were analyzed using pLink software and manually reviewed to detect the disulfide linkage between PorG and PorK.

To identify glycosylation sites on PorK and PorN, the purified PorKNG complex was electrophoresed on an SDS-PAGE gel. The bands corresponding to PorK and PorN were subjected to trypsin digestion and the tryptic fragments analysed as essentially described in [40]. Proteins and peptides were identified using Mascot software. All searches were conducted using trypsin with up to two missed cleavages allowed, peptide mass tolerance was set to 10 ppm, fragment mass tolerance to 0.04 Da, and a fixed modification of carbamidomethyl (C), variable modifications of oxidation (M), 792.25 (S, T), 1352.42 (S, T), 1394.42 (S, T) and 1436.43 (S, T).

### Gene cluster analysis

Gene cluster analysis was performed using the Kegg SSDB gene cluster tool, accessible from https://www.genome.jp/kegg/genes.html, using default parameters (gap size = 0; Threshold = 100).

### AlphaFold modelling and dissociation constant (K_d_) calculations

Model structures for various components of T9SS and their complexes were generated using AlphaFold3 through the AlphaFold3 server interface [41]. Where necessary membrane effects on proteins during model building were simulated by the inclusion of multiple copies (usually 100) of myristic acid in the modelling process. Full membrane complexes were generated for protein models using the OPM [42] and CHARMM-GUI [43, 44] servers. Interfacial energies in protein-protein complexes were predicted using the PRODIGY server [45, 46] using default settings other than the temperature, which was set to 37°C.

## Results

### High-resolution structure of the PorKN rings

Previously, we purified the PorKNG complex from *P. gingivalis* and demonstrated that it forms large, 50 nm ring-shaped structures [19]. However, due to one preferred orientation on the EM grid and protein aggregation, we were only able to achieve a resolution of ∼40 Å. At this resolution, we could discern that the structure was composed of one large ring and one slightly smaller ring, but the organization of the PorK, PorN and PorG subunits within the ring structure remained unclear. With optimization of buffer conditions, which included addition of 1M urea prior to applying the sample on an EM grid, we were able to mitigate strong aggregation of the PorKN rings and capture various orientations of the rings on the EM grid by focusing acquisition on thicker vitreous ice. This improvement enabled us to achieve ∼3.5Å maps of the PorKN rings (**Fig 1B**, 1C and 1D, Extended Data Fig 1). Several rounds of classifications were performed that resulted in two classes with distinct symmetry of C32 (46% of final particles) and C33 (54% of final particles). A very small number of top views with C34 and C35 fold-symmetry were observed early in the 2D classification process but did not lead to 3D classes suggesting that these two species are too rare to bear on the classification. *P. gingivalis* is therefore capable of assembling rings of different stoichiometry. The interfaces between adjacent heterodimers were identical in the C32 and C33 fold symmetries, however, they did make a difference to the measured diameter of the ring: an increase of 16 Å was observed in the C33 fold symmetry (529 Å) compared to the C32 fold-symmetry (513 Å). In both instance each heterodimer had the periodicity of 50.36 Å.

The AlphaFold [41] predicted structure of the PorK and PorN heterodimer was used as a starting point for structure refinement. The structure of PorK was solved from Glu^45^ to Ala^481^ missing the N-terminal 21 residues (not including signal peptide) and the C-terminal 10 residues. PorN was solved from Leu^50^ to Gln^297^ missing the N-terminal 25 residues (not including signal peptide) and the C-terminal 63 residues. Although present in the purified samples, PorG was unable to be resolved (**Fig 4B**) [19]. The orientation of the ring was easily solved by comparison to the published in-situ structures of the ring where the larger diameter “upper ring” is adjacent to the OM [21]. The top and side views of the ring show that PorK forms the upper ring, featuring a large, flat surface orientated towards the OM that is bound by a discontinuous “outer rim” (**Fig 1C and 1D**). PorN forms the second ring of a slightly smaller external diameter underneath PorK and it folds to create cog-like projections that *in vivo* would be on the periplasmic side (**Fig 1C**). Enlarged side-views show how PorK subunits interlock through a large wedge-shaped protrusion, which extends diagonally down and across to fit into a crevice on the adjacent monomer (**Fig 1E and 1G**). Interestingly, the outside surface of the PorK outer rim resembled a three-toed claw that exhibited highly conserved negatively charged residues (**Extended Data** Fig 2A and 2B). In *P. gingivalis* PorK, these were largely contributed by Glu^121-2^, Glu^127^, and Glu^152-4^. A small N-terminal region of PorK (residues Gly^50^-Val^103^) and the C-terminal part of PorK (residues Trp^261^-Arg^471^) together adopt the fold of a formylglycine generating enzyme (FGE) which makes up most of the core of PorK and a large part of the cavity that binds PorN (**Extended Data** Fig 2C). In contrast, the region Arg^104^-Cys^260^ is unique to PorK and includes the interlocking wedge, the negatively charged outer rim and the outside wall of the PorN-binding cavity (**Extended Data** Fig 2C).

A cross-section view of the PorKN heterodimer (**Fig 1F**) highlights the location of the most N- and C-terminal parts solved. For PorK, both appear on the OM side, consistent with the N-terminal Cys being lipidated and inserted into the membrane. The most C-terminal part of PorN was the α-helix extending from Thr^271^-Arg^290^ located on the inside of the ring extending toward the OM (**Fig 1F**). This helix can be seen in the inside views appearing as vertical bars (**Fig 1C**). The unsolved N- and C-terminal regions of PorK and PorN are likely to be flexible in the purified rings. The unresolved C-terminal region of PorN (residues Thr^291^-Lys^361^) is highly positively charged and is predicted to be mostly α-helical.

PorN was found to insert into a cavity produced underneath each pair of PorK subunits (**Fig 1G**). Three linear regions of PorN are involved in the binding. The first region from Leu^50^ to Trp^70^ is mostly α-helix and binds deeply within the cavity of PorK (**Extended Data 3A**). Within this region, Arg^53^ and Ser/Thr^51^ are absolutely conserved among 45 diverse PorN homologs within the *Bacteroidia* class. These residues penetrate deeply into a pocket of PorK and are modelled to interact mainly through hydrogen bonds (**Extended Data 3A**). The second and third regions from Phe^179^ to Ser^184^ and Glu^285^ to Val^295^ respectively form one deeply penetrating loop, and several residues involved in hydrophobic and aromatic stacking interactions (**Extended Data 3A**). Together, the nature and extent of these interactions suggest a stable interaction between PorK and PorN. Consistent with this, the estimated dissociation constant (K_d_) of one PorN subunit binding to two PorK subunits at 37°C was computed to be 2x10^-11^ M with a ΔG of -15.2 kcal/mol, indicative of a strong association.

The cog-like projections of PorN suggest potential interaction points with the dimeric PorM rotor. We therefore predicted the binding site for this known interaction using AlphaFold. The C-terminal d4 domain of PorM was predicted to bind between the cog-like projections of PorN, potentially through electrostatic interactions between the conserved negatively charged cogs and the positively charged rotor tip (**Extended Data** Fig 3B). While the ipTM score was 0.47, the prediction was consistent, and the geometry was a good fit. The poor score may reflect that the binding is dynamic-the main requirement being that the PorM shaft can slot in-between the cogs that can drive rotation of the rings.

### O-glycosylation of PorK and PorN

PorK and PorN were predicted to be *O*-glycosylated at three and two sites respectively, on the basis of the published (D)(S/T)(A/I/L/V/M/T/S/C/F/G) glycosylation motif [40]. We performed mass spectrometry analysis on the purified PorK and PorN protein bands resulting in the identification of glycosylated peptides that covered all 5 sites (**Table 1**). All sites were modified with the known major heptasaccharide O-glycans of Δmass 1436 and 1394 Da, while the minor O-glycans of Δmass 1352 and 792 were observed for only some of the sites. The level of site occupancy was high since non-glycosylated forms of the same peptides were not observed, with the exception of non-glycosylated PorN ^264^ALAEYCPTPDSMK^280^ which was detected at a low level. The collision-induced decay (CID) MS2 spectra clearly show the presence of the heptasaccharide associated with each of the five glycosylated peptides (**Extended Data** Fig 4).

**Table 1:**
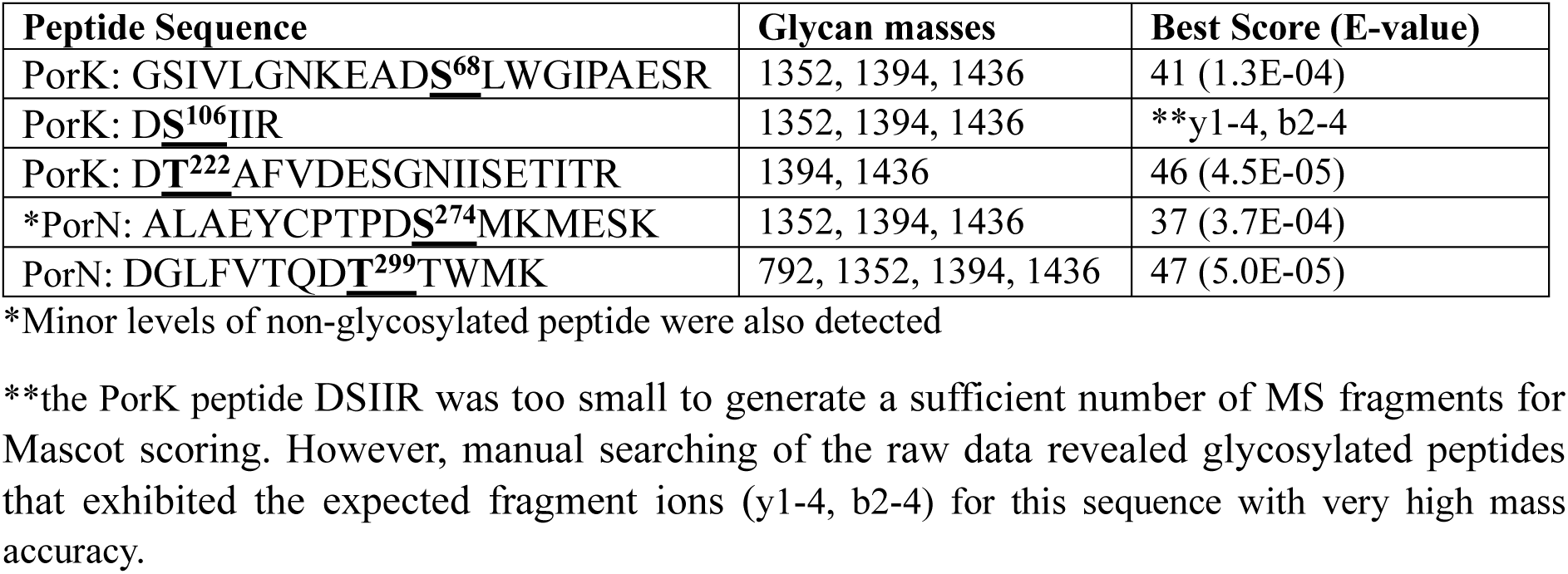
Glycosylated peptides of PorK and PorN.

The PorK glycans were also observed in the cryo-EM map. PorK Ser^68^ was observed with only one sugar, while PorK Ser^106^ and Thr^222^ were observed with full-length (heptasaccharide) glycans with only the acetyl glycerol phosphate portion unassigned (**Fig 1H**). The glycans on Ser^106^ and Thr^222^ are located in the PorK-specific region under the outer rim with the terminal sugars potentially interacting with adjacent PorK subunits. In contrast, the glycans on PorN were not visualised on the cryo-EM map, suggesting high flexibility.

### Role of PorG in the T9SS

To investigate the role of PorG within the PorKN rings and the T9SS, we generated a *porG* deletion mutant in the *P. gingivalis* W50 strain. We assessed the impact of the deletion on T9SS function through phenotypic analysis of pigmentation, gingipain activity assay and the ability of T9SS cargo proteins to attach to the cell surface. The W50 *porG* mutant was apigmented on blood agar plates as opposed to the black pigmented WT, consistent with earlier reports of a *porG* deletion mutant in the *P. gingivalis* ATCC 33277 strain [15]. This suggests that PorG is an essential component of the T9SS (**Fig 2A**). The *porG* mutant displayed negligible <5% cell surface and supernatant gingipain (Kgp and Rgp) activity compared to the WT (**Fig 2B**), consistent with an inability of these T9SS dependent proteases to be translocated across the OM. In contrast to the WT, anti-A-LPS western blot showed no signal above 60 kDa in the *porG* mutant (**Fig 2C**), indicating the inability of T9SS cargo proteins to be attached to the cell surface by modification with A-LPS. Accordingly, the layer of cargo proteins on the cell surface known as the EDSL was absent in cryo-electron microscopy images of *porG* mutant cells but present in the WT (**Fig 2D**). We also constructed *porG* complementation strains (*porG*^+^), by chromosomal integration of *porG* into the *fimA* (*pg2132*) locus. The *porG*-complemented strain demonstrated pigmentation and increased protease activities (**Fig 2A and 2B**), indicating the phenotype observed in the *porG* mutant is specifically due to the lack of PorG. Jointly, these results indicate that PorG is an essential component of the T9SS.

**Figure 2:**
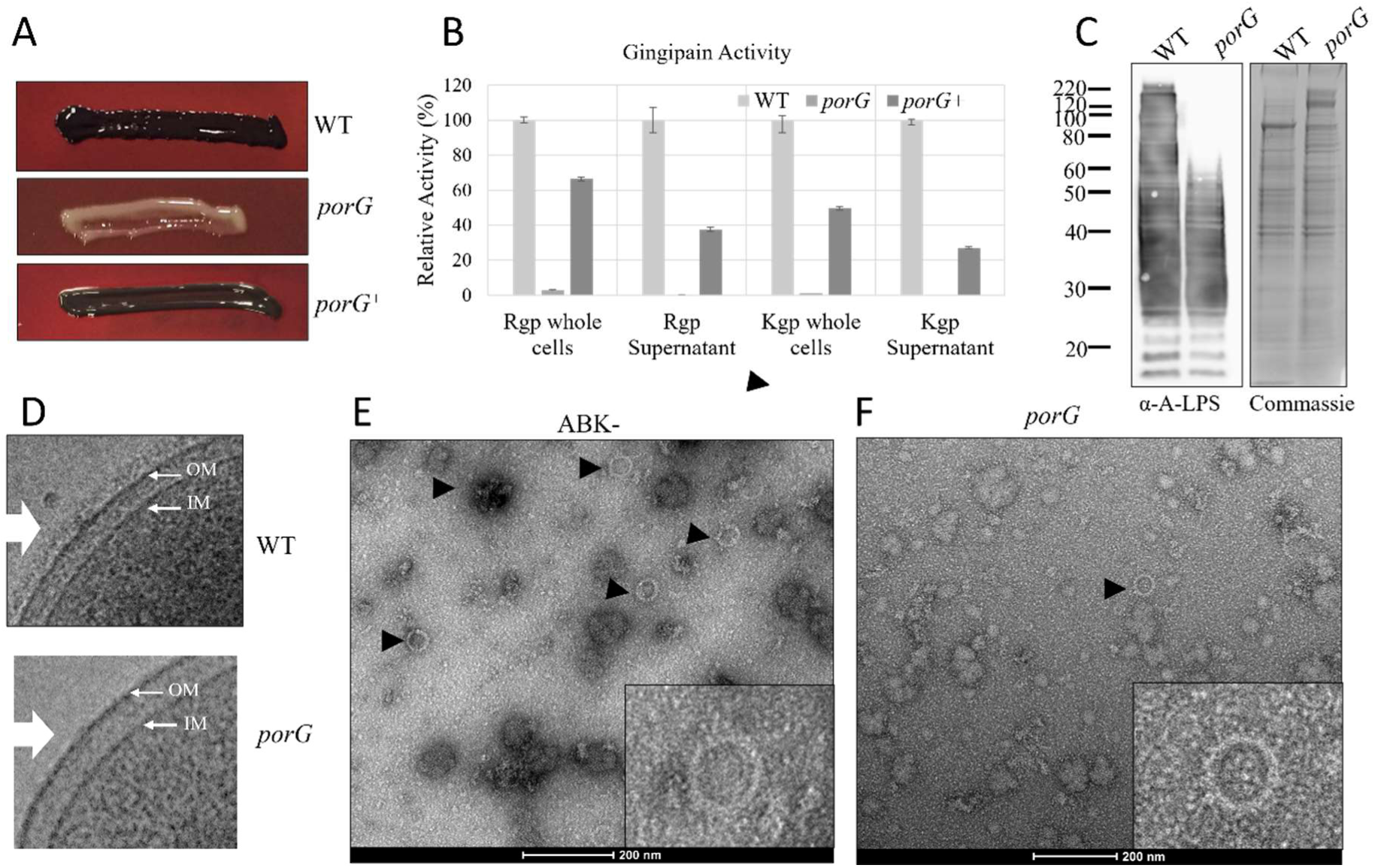
PorG is an essential component of the T9SS. (A) *P. gingivalis* WT, *porG* deletant and *porG^+^* complementation strain pigmentation on blood agar plates after 7 days of incubation. (B) Intact whole cells and supernatant samples were analyzed for endoproteolytic activities of the T9SS RgpA, RgpB and Kgp cargo proteins using Arg- and Lys-specific chromogenic substrates. Relative gingipain activities are represented as a percentage of the WT activity for all fractions and strains. (C) Cell lysates from WT and the porG mutant were electrophoresed using SDS-PAGE and the proteins were transferred onto a nitrocellulose membrane and probed with mouse monoclonal antibody mAb 1B5 (anti-A-LPS). (D) Cryo-electron micrographs showing the presence of an electron-dense surface layer (*EDSL*) in WT cells, and the absence of this layer in the *porG* mutant cells (white arrows). Arrows show outer membrane (OM) and inner membrane (IM). (E, F) Large protein complexes were prepared from (E) ABK- and (F) *porG* mutant cells, the samples were negatively stained and viewed under the electron microscope. The electron micrographs show the presence of large (50 nm) rings (black arrows). Scale bars, 200 nm.

Next, we investigated if the PorKN rings were affected by the absence of PorG. We applied our partial purification protocol for macromolecular complexes, that we have previously used to assess the presence of the rings in various T9SS mutants [19]. For comparison with the porG mutant, we used the *P. gingivalis* ABK-strain which lacks all three gingipains but has a functional T9SS. The samples were stained with uranyl acetate and visualised under electron microscopy. The 50 nm PorKN rings were visible in the ABK-cells (**Fig 2E**) and the *porG* deletion mutant (**Fig 2F**).

### The *porG* gene is located within a conserved gene cluster

*P. gingivalis porG* (*pg0189*) is located adjacent to genes coding a BamA-associated lipoprotein (PG0188) [47] and undecaprenyl diphosphate synthase (UppS, IspU, PG0190). Nearby are also the genes coding BamA (PG0191), and two Skp proteins (PG0192 and PG0193) [24]. This six-gene cluster was found to be conserved across numerous genera in the *Bacteroidota* phylum, especially in class *Bacteroidia*, but also extending to other classes such as *Cytophagia*, *Sphingobacteriia* and *Flavobacteriia* with the exception of the lipoprotein orthologs which were sometimes undetected (**Supplementary table 2**). Besides UppS which is involved in peptidoglycan and LPS biosynthesis, the other four encoded proteins are involved in Omp biogenesis [48], with Skp functioning as a periplasmic chaperone and the BAM complex responsible for the insertion of Omps into the OM [49]. The conserved location of *porG* within this cluster therefore suggested a possible involvement with Omp biogenesis that is specific to the T9SS. Some species of *Bacteroides* such as *B. fragilis*, while closely related to *P. gingivalis,* lack PorG and the T9SS. A gene cluster analysis of *B. fragilis bamA* (BF0504), shows an extended gene cluster that includes all the genes noted above in the same order except for *porG* (**Supplementary table 2**).

### PorG is not required for the correct localisation of the T9SS OM β-barrel proteins

To explore if the T9SS components are affected in the absence of PorG, we performed label free quantitative (LFQ) proteomics on the whole cell lysate of WT and the *porG* mutant as per our published protocol [50]. In the *porG* mutant all the T9SS components were decreased in abundance compared to the WT (**Extended data Fig 5**), including 6 OM β-barrels, however OM β-barrels unrelated to the T9SS were unaffected. This suggest that the T9SS is downregulated in the absence of PorG.

BamA is a highly conserved protein responsible for folding and inserting OM β-barrel protein into the OM [51]. Since the *porG* gene (*pg0189*) is localised near the *bamA* gene (*pg0191*), we explored whether PorG was involved in the localisation and folding of the T9SS specific OM β-barrels Sov, PorV, PorQ, PorT, PorF and PorP. The cell envelope (CE) proteins from both the wild-type (WT) and *porG* mutant strains was subjected to trypsin digestion and analyzed by mass spectrometry. The data were analysed by MaxQuant software, and the values of the ratio of LFQ intensities of the *porG* mutant to WT was normalised to other OM β-barrels that are not related to the T9SS. The T9SS OM β-barrels were identified in the CE of the *porG* mutant albeit at reduced levels (**Fig 3A**), consistent with the whole cell lysate results (**Extended Data** Fig 5). Together, this suggests that T9SS OM β-barrels are able to integrate into the OM in the absence of PorG. We also investigated the protein interactions of the Sov translocon in the *porG* mutant by blue native PAGE western blot and affinity purification mass spectrometry using Sov specific antibodies. In the native western blot, two Sov specific bands were observed, which we previously assigned to mainly PorV-Sov (∼500 kDa) and PorV-Sov-PorW (∼750 kDa) (**Fig 3B**) [23] consistent with the PorV and Sov β-barrels being properly integrated in the OM. Of note, to obtain the signal in the *porG* lane, the blot was exposed for longer supporting the finding of lower Sov abundance in the *porG* mutant. Similarly, we previously employed Sov co-immunoprecipitation studies in several strains to show that the Sov interactome includes PorA, PorD, Plug, PorK and PorN in addition to PorV and PorW [23]. To test if Sov can make the same interactions in the *porG* mutant, the Sov affinity enriched material from ATCC 33277 *porG* mutant (gift from Prof Naito) was analysed by mass spectrometry and found to contain Sov, PorV, PorA, PorW and PorK at more than 10 times the level observed in the negative control while PorD was also present but its observed ratio was not significant (**Fig 3C**). The ATCC 33277 *porG* mutant was selected for this work to maintain consistency with our previous published work in the ATCCC 33277 *P. gingivalis* strain [23]. Taken together, these results strongly suggest that the two key T9SS Omps, Sov and PorV can integrate into the OM and interact with their binding partners suggesting that the defect in the T9SS function due to lack of PorG is not due to the inability of Omps to be incorporated into the OM.

**Figure 3:**
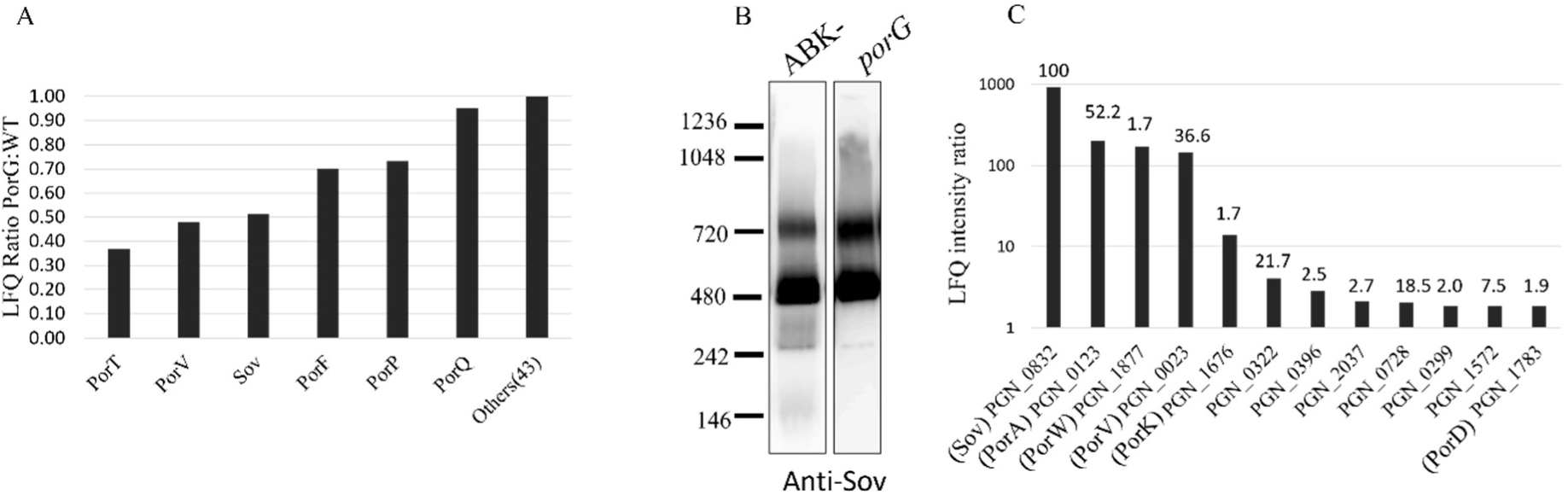
PorG is not involved in localising T9SS OM β-barrels to the CE. (A) Cell envelope fractions from WT and the *porG* mutant were analysed by mass spectrometry and MaxQuant software. The LFQ ratios of *porG* mutant to WT of the T9SS OM β-barrels were plotted relative to other Omps not related to the T9SS. (B) Cell lysates from ABK- and the *porG* mutant were electrophoresed on a blue native gel. The proteins were transferred onto a PVDF membrane and probed with anti-Sov antibodies. (C) Cell lysates from *porG* and *sov* mutants were subjected to co-immunoprecipitation using anti-Sov antibodies. The enriched material was analysed by mass spectrometry and MaxQuant software. The graph is a plot of LFQ intensities ratio of *porG* to *sov* mutant (log scale). The values of abundance of each protein relative to Sov (iBAQ) is written on top of the bar graph.

### PorG forms a disulfide bond with PorK

To further understand the nature of the interaction between PorK, PorN, and PorG, we used AlphaFold Multimer [41] to predict the structure of the PorKNG complex and its relationship with the lipid bilayer. The modelling revealed that PorK anchors to the membrane via its lipid moiety and interacts with PorG, while also showing a gap of ∼ 20 Å between the flat surface of PorK and the OM (**Fig 4A**). Assuming a conventional OM β-barrel architecture where the N-and C-termini are located in the periplasm, the model shows the periplasmic end of the PorG barrel interacting with PorK (**Fig 4A**). The PorG C-terminal extension containing Cys^232^ was predicted to run on the surface of PorK with the Cys placed in close proximity to PorK Cys^356^, suggesting potential for disulfide bond formation (**Fig 4A**). The structure of PorKN derived from the cryo-EM analysis confirmed the location of the PorK Cys^356^ and its availability for creating a disulfide bond with PorG.

To verify the presence of a disulfide bond between PorK and PorG, the purified PorKNG complex sample was analysed by SDS-PAGE under denatured reducing and non-reducing conditions (**Fig 4B**). There were no obvious differences in the migration profiles of the two prominent protein bands of PorK and PorN under reducing and non-reducing conditions. However, under non-reducing condition the intensity of the PorG (∼25 kDa) (band 4) (**Fig 4B**) decreased significantly and a band at ∼90 kDa appeared (band 1) (**Fig 4B**). This suggested the presence of a disulfide bond between PorG and its associated partner. To investigate which proteins are present in the ∼90 kDa band, this protein band together with the PorK and PorN bands in the non-reducing gel lane were excised and cut in half. Half of each band was subjected to trypsin digestion in the absence of DTT while the other half was reduced and alkylated before trypsin digestion. The tryptic fragments were analysed by mass spectrometry.

**Figure 4:**
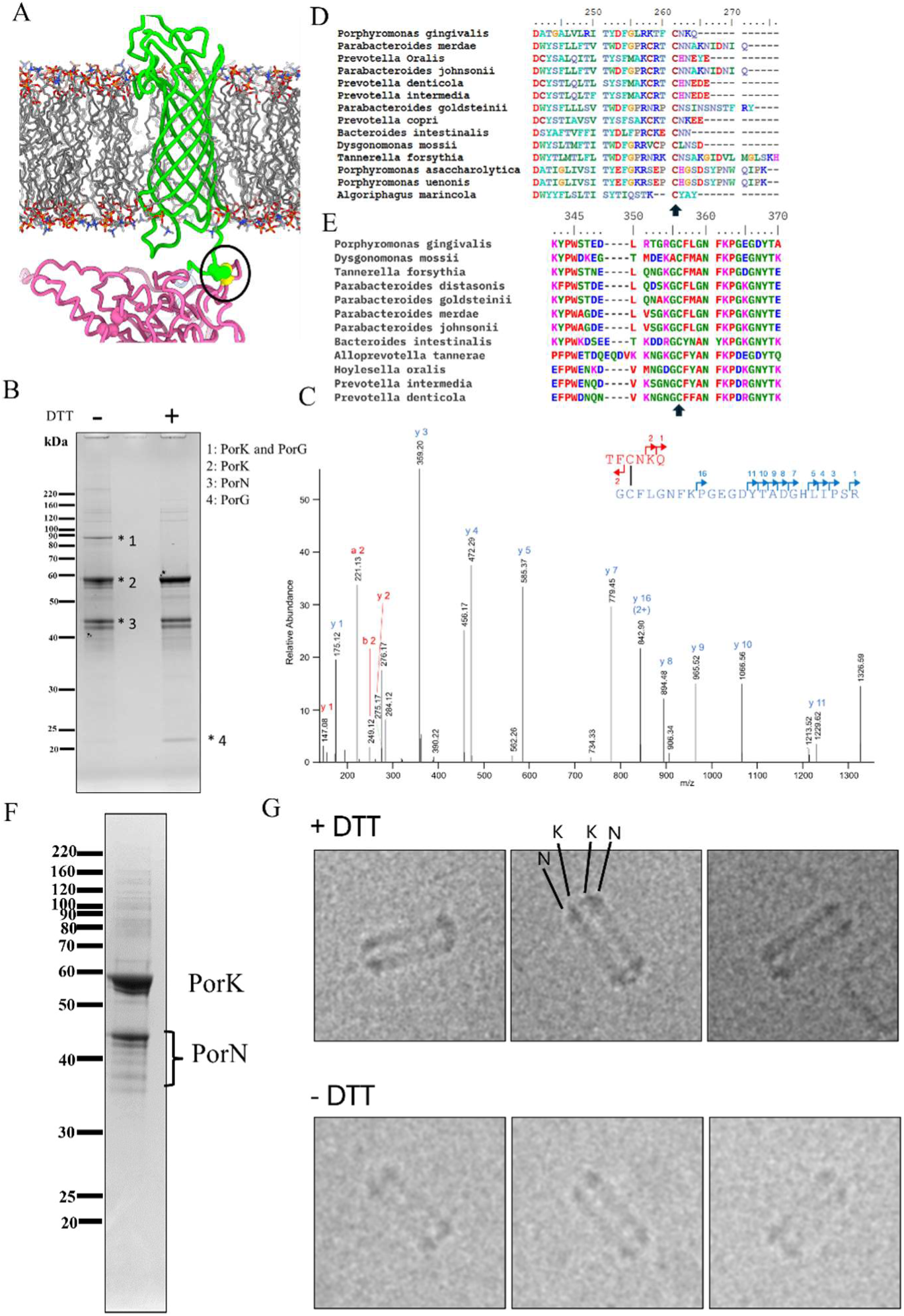
Conserved Cys^232^ of PorG forms a disulfide bond with a conserved Cys^356^ of PorK. (A) AlphaFold3 model of PorG (green)-PorK(pink) interaction showing the close proximity of the PorG Cys^232^ and PorK Cys^356^ (circled). Cys residues are shown in ball rendering (B) SDS-PAGE of purified PorKNG complex electrophoresed under reducing (DTT +) and non-reducing (DTT -) conditions. (C) MS/MS spectra consistent with a disulfide bond peptide between PorG (^230^TFCNKQ^235^) and PorK (^355^GCFLGNFKPGEGDYTADGHLIPSR^378^). (D) Alignment of the C-termini of representative proteins identified by BLAST search to be homologous to PorG. The conserved cysteine is indicated by the arrow. (E) Alignment of residues of representative proteins identified by BLAST search to be homologous to PorK. The conserved cysteine is indicated by the arrow (F) SDS-PAGE of purified PorKNG complex under reducing conditions. (G) Cryo-EM images of the PorKN rings showing the purified dual ring structure (NK-KN) in the presence (top panel) and absence of DTT (bottom panel).

Analysis of the band at ∼90 kDa identified both PorK and PorG proteins, however, pLink software did not identify any disulfide bonded peptides. Thus, the MS/MS spectra were analysed manually to identify the disulfide bond. PorG only contains one cysteine, Cys^232^. In the reduced gel band analysis of the ∼90 kDa band a PorK peptide containing Cys^356^ was identified “^355^GCFLGNFKPGEGDYTADGHLIPSR^378^”. Of note this peptide was absent in the non-reduced gel analysis of the same band. Thus, we created a table of expected masses containing the PorK peptide “^355^GCFLGNFKPGEGDYTADGHLIPSR^378^” combined with four differentially cleaved PorG peptides “^230^TFCNK^234^”, “^230^TFCNKQ^235^”, “^229^KTFCNK^234^” and “^229^KTFCNKQ^235^” (**Table 2**). Analysis of the MS1-level spectra accurately identified MS peaks corresponding to the PorK peptide cross-linked to all four of the variant PorG peptides (**Table 2**). Each of these peaks was only found in the non-reduced band supporting their identity as crosslinked peptides. The assignments were confirmed from multiple MS/MS spectra, from various charge-states with the best spectrum obtained for the crosslinked peptides PorK- ^355^GCFLGNFKPGEGDYTADGHLIPSR^378^ and PorG-^229^KTFCNKQ^235^ with a parent ion mass of 822.8873 (4+) (**Fig 4C**). Bioinformatic analysis and BLAST searches of PorG and PorK identified PorG, Cys^232^ and PorK Cys^356^ to be conserved suggesting that the crosslink between these two Cys residues may be a conserved and important feature (**Fig 4D and 4E**). Since PorG was not resolved in the high-resolution structure of the PorKN rings we determined its stoichiometry by densitometry analysis. Densitometry analysis of coomassie-stained gels of the purified complex estimated ∼6-7 PorG molecules per PorKN ring (**Extended Data** Fig 6).

**Table 2:** Expected and observed masses (M/Z) of PorK peptide (GCFLGNFKPGEGDYTADGHLIPSR) + PorG peptides.

| PorK peptide crosslinked with: | Calculated m/z | Observed m/z non-reduced | Observed m/z reduced |
| --- | --- | --- | --- |
| PorG: TFCNK | 790.8714 (4+) | 790.8713 (4+) | Not observed |
| PorG: TFCNKQ | 822.8860 (4+) | 822.8873 (4+) | Not observed |
| PorG: KTFCNK | 822.8951 (4+) | 822.8946 (4+) | Not observed |
| PorG: KTFCNKQ | 854.9098 (4+) | 854.9090 (4+) | Not observed |

To investigate a structural role of disulfide bond formation between PorK and PorG, we employed our PorKNG complex purification protocol using reducing conditions at all steps and electrophoresed the purified sample on an SDS-PAGE gel. We identified the PorK and PorN bands, however, the PorG band was not detected (**Fig 4F**). PorN appeared to be degraded slightly, perhaps due to some protease activity. The PorKN containing complex isolated under reducing condition was then visualised under cryo-EM to investigate the effect of reduction and the removal of PorG on the structure of the rings. The rings under reducing conditions appeared to be same as the rings without DTT (**Fig 4G**), and the dual ring structure (NK-KN) was also unaffected. This suggests that once the rings have formed, the removal of PorG does not impact the PorKN ring structure. In the presence of DTT, the PorG is removed but the dual ring structure stays intact likely due to the lipid interactions of the PorK lipoprotein. In the absence of the DTT PorG is likely to be present in between the lipids and due to its low stoichiometry, are difficult to visualise. This suggests that the artefact of the dual ring formation between the rings is attributable to the lipids in PorK as we had speculated previously [19] and is not related to the presence of PorG.

### The PorG-PorK disulfide bond is essential for the function of the T9SS and is not required for the formation of stable PorKN rings

To further understand the role of the disulfide bond between PorG and PorK, we mutated PorG Cys^232^ to Ser^232^. We mutated the Cys^232^ codon of *porG* with insertion of *ermF* 3’ as a recombinant marker gene producing the strain designated PorG^C232S^. As a control, a recombinant cassette in which the Cys^232^ codon was unchanged was also prepared, this strain was named PorG^Cys-Cys^. The control PorG^Cys-Cys^ strain retained a WT like black pigmentation on blood agar indicating the recombination strategy did not affect the function of the T9SS (**Fig 5A**). In contrast, the PorG^C232S^ strain was similar to the *porG* mutant in that it was not pigmented on blood agar (**Fig 5A**), had no A-LPS signal above 60 kDa in the mAb-1B5 western blot (**Fig 5B**), and had no EDSL layer (**Fig 5C**). All these findings indicate that the T9SS is non-functional in the PorG^C232S^ strain. To explore if the PorK and PorN were localised to the CE, we prepared CE fractions from WT and PorG^C232S^ mutant. The sample was subjected to western blot analysis using PorK and PorN antibodies. The immunoblot showed the presence of both PorK and PorN with similar abundances in both WT and PorG^C232S^ mutant (**Fig 5D**). Next, we investigated if the PorKN rings were formed in the PorG^C232S^ mutant by isolating the complex using our published purification protocol under non-reducing conditions [19]. The sample was analysed on the gel to assess its purity and the identity of the proteins present (**Fig 5E**). Both PorK and PorN bands were identified, however, the PorG band was missing on the gel. This suggested that the disulfide bond between PorG and PorK is required for the stable interaction. The sample was then negatively stained and visualised under the electron microscope. The rings were visualised and appeared to be of unaltered size and shape (**Fig 5F**), thus Cys^232^ of PorG is not essential for PorKN ring formation. Jointly, these results suggest that PorG may play an active role in the mechanism of type IX secretion that is dependent on its disulfide bond to PorK.

**Figure 5:**
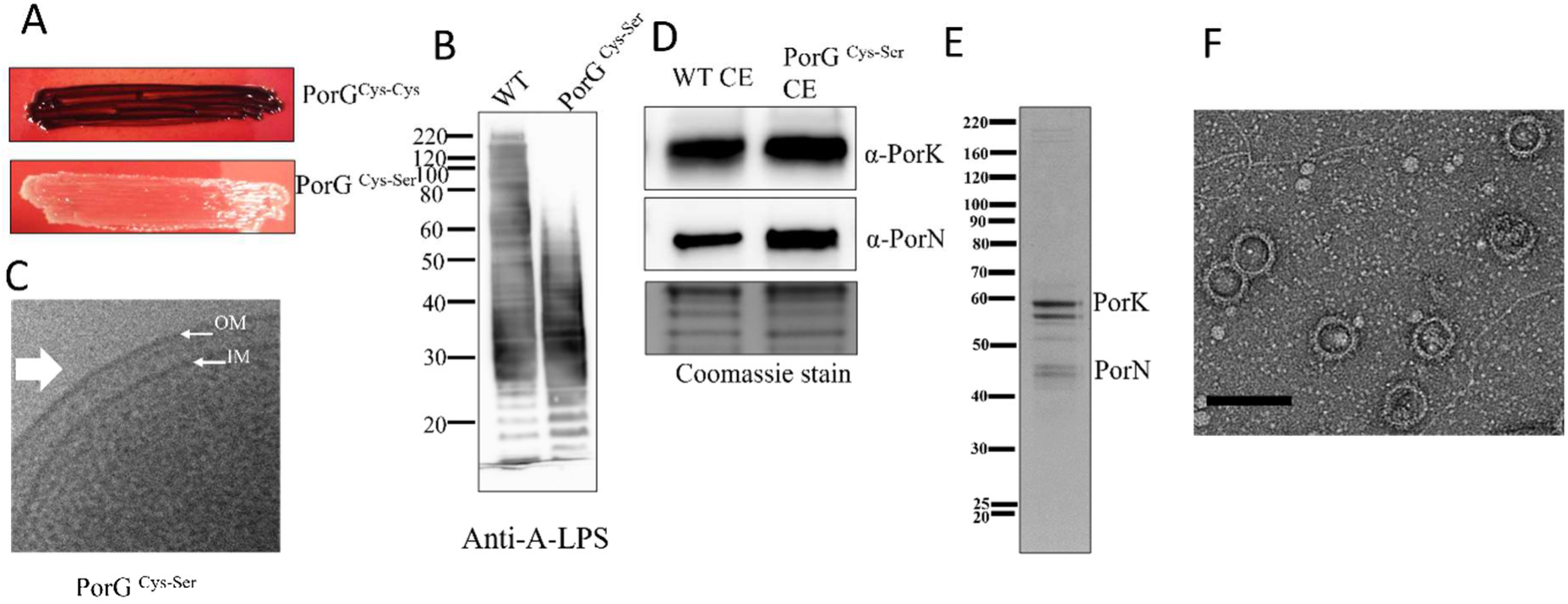
The disulfide bond between PorG and PorK has a functional role in the T9SS. (A) *P. gingivalis* PorG^Cys232-Cys^, PorG^Cys232-Ser^ pigmentation on blood agar plates after 7 days of incubation. (B) Cell lysates from WT and the PorG^Cys-Ser^ were electrophoresed on an SDS-PAGE gel and the proteins were transferred onto a nitrocellulose membrane and probed with mouse monoclonal antibody mAb 1B5 (anti-A-LPS). (C) Cryo-electron micrographs showing the absence of an EDSL (white arrow) on the PorG^Cys-Ser^ mutant cells. (D) Cell envelope fractions from the WT and PorG^Cys-Ser^ were electrophoresed on an SDS-PAGE gel. The proteins were transferred onto a nitrocellulose membrane and probed with PorK and PorN antibodies. The Coomassie stained gel shows the protein load. (E) SDS-PAGE gel of purified PorKN complex from the PorG^Cys232-Ser^ mutant cells. The protein band observed beneath PorK was identified as PG1881, which co-purified with the complex. (F) Purified PorKN complex from PorG^Cys-Ser^ was negatively stained and visualised by cryo-electron microscopy, showing ring structures. Scale bar: 100 nm.

### The Attachment Complex associates with the PorKN rings

Although the function of the Attachment Complex comprising PorU, PorV, PorQ and PorZ is known, its structure and organization relative to other T9SS sub-complexes is undiscovered. Therefore, we investigated whether the Attachment Complexes were associated with the PorKN rings. To address this question, we performed co-immunoprecipitation (Co-IP) with PorZ antibodies using lysates from ABK-, *porV*, *porU*, and *wbaP* mutant *P. gingivalis* cells. The immunoprecipitated material was quantified by mass spectrometry and MaxQuant relative to *porZ* mutant negative control. The label-free quantitation (LFQ) intensity ratios of the top 30 proteins in the ABK Co-IP were plotted on a heatmap, along with the LFQ intensity ratios for the *porV*, *porU*, and *wbaP* mutants (**Fig 6A**). As expected, the other attachment complex components PorQ, PorU and PorV were observed in the pulldown assay with high LFQ intensity ratios (**Fig 6A**). PorK, PorN, PorG, PorE, PorT and Sov also exhibited high LFQ intensity ratios (**Fig 6A**), indicating that their presence in the pulldown assay was specific to the presence of PorZ. The iBAQ metric was used to quantify the relative abundance of proteins in the pulldown. After correction of the iBAQ values using the *porZ* mutant negative control, the values were quantified relative to PorQ. PorZ, PorU, PorQ and PorV were the most abundant proteins present in the immunoprecipitated material followed by PorK, PorN and PorG consistent with the Attachment Complexes being associated with the PorKN rings and PorG (**Fig 6B**). The Co-IP using PorZ antibodies was also performed on *porV*, *porU* and *wbaP* mutant cells (**Table 3**). In the *porV* and *porU* mutants, PorU and PorV proteins were absent in the immunoprecipitated material as expected, but the PorKN rings and PorG were still present (**Table 3**), suggesting that the PorQZ subcomplex [25, 32] can form in the absence of PorU and PorV and still associate with the PorKN rings and PorG. A-LPS binds to PorZ and is a substrate of PorU [26]. In the *wbaP* mutant which does not express A-LPS, all four components of the attachment complex were enriched together with the components of the PorKN rings and PorG (**Table 3**), indicating that A-LPS was not required for their association. Collectively, this suggests that the Attachment Complex associates directly with the PorKN ring and PorG.

**Figure 6:**
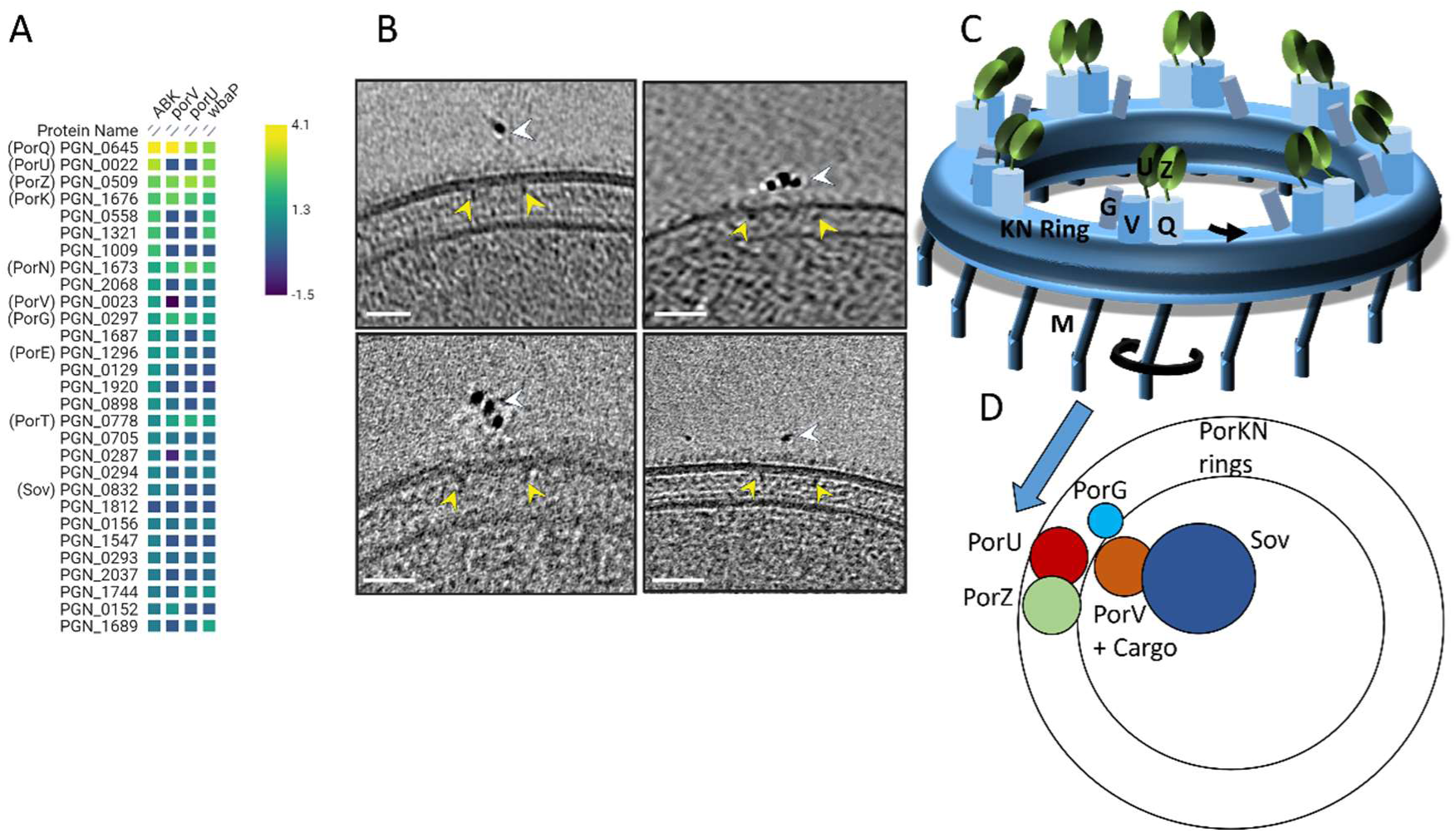
Association of the attachment complexes with the PorKN rings. The *P. gingivalis* strain lysates were subjected to coimmunoprecipitation using PorZ-specific antibodies. The *porZ* mutant was used as a negative control. The immunoprecipitated samples were digested with trypsin, analyzed by mass spectrometry, and quantified using MaxQuant software. The ratio of LFQ intensity relative to the *porZ* control was determined. (A) The top 30 proteins for (ABK^−^/*porZ*) were plotted on a heatmap along with LFQ intensity ratios for the same proteins calculated for *porV*, *porU* and *wbaP* mutants. (B) The *porV* mutant cells were labelled with PorZ primary antibodies followed by anti-rabbit gold (6 nm) secondary antibodies. The *porZ* mutant cells were used as a negative control. Cryo-ET was performed on the immunolabeled cells. In cryotomograms, PorKN rings were observed directly underneath the gold clusters. White arrows point to gold clusters and the yellow arrows point to the PorKN rings. (C) A cartoon representation of the PorKN rings connected to PorM (M) on the periplasmic side, and to the outer membrane (OM) β-barrels, including PorG (G), PorQ-PorZ (Q, Z), and PorV-PorU (V, U) on the OM side. (D) A schematic illustrating the proposed mechanism of the T9SS secretion and the attachment of cargo proteins to the cell surface. The rotary movement of PorM causes the PorKN ring and its’ associated proteins (PorG and attachment complexes) to rotate along with it. The Sov-PorV-cargo is positioned adjacent to the PorKN rings, held in place by the transient interaction between PorW and PorN. As PorG rotates, it provides the energy to push the PorV-cargo out of the Sov translocon and place it next to the attachment complex for sortase (PorU) reaction and anchorage to the cell surface via A-LPS.

**Table 3:** Relative abundance of PorZ binding partners in various T9SS mutants.

| Protein Name | LFQ Ratio ABK-/porZ | *iBAQ (%)* |  |  |  |
| --- | --- | --- | --- | --- | --- |
|  |  | ABK- | *porV* | *porU* | *wbaP* |
| PorZ | 655 | 255 | 324.8 | 389 | 679 |
| PorQ | 11810 | 100 | 100 | 100 | 100 |
| PorU | 3003 | 70.8 | 0 | 0 | 93 |
| PorV | 43 | 91.7 | 0 | 0 | 80 |
| PorK | 327 | 7.7 | 12.4 | 16.2 | 25 |
| PorN | 67 | 4.6 | 7.8 | 12.8 | 25 |
| PorG | 42 | 0.4 | 1.4 | 1.2 | 2 |
| PorT | 16 | 0.1 | 0.5 | 0.3 | 0.4 |
| PorE | 28 | 0.1 | 0.1 | 11.9 | 0 |
| Sov | 10 | 0 | 0 | 0 | 0 |

To validate this finding and investigate if multiple Attachment Complexes were co-located and are above the PorKN rings *in-vivo*, *porV* mutant cells were labelled with anti-PorZ and secondary antibodies conjugated to gold and imaged by cryo-electron tomography (cryo-ET). The *porV* mutant was selected for this work because it lacks an EDSL, providing better access of the antibodies to the PorZ antigen. From 62 tomograms of *porV^-^* cells we observed an average of 4 clusters of gold particles per cell. Within each cluster were up to 4 gold particles and tomogram sections showed the PorKN rings directly beneath the gold clusters (**Fig 6B**). In comparison, nanogold cryo-ET on a *porZ* mutant only showed a small number of labelled cells relative to the *porV* mutant, indicating the specificity of the PorZ antibodies. Together, these results indicate that multiple Attachment Complexes are co-located directly above the PorKN rings.

## Discussion

The T9SS in *P. gingivalis* is comprised of at least 20 proteins. We previously showed that the PorK and PorN components of the T9SS form large 50 nm ring-shaped structures which are intimately associated with PorG [19]. The exact role of these rings in the mechanistic function of the T9SS is unknown. We proposed that the large PorKN rings create a diffusion barrier that retains all the T9SS components within the ring to promote efficient and coordinated protein secretion and cell surface attachment [52]. The PorKN ring makes a direct contact with the PorL/M motor [21] that powers the T9SS therefore the ring is also presumably involved in distributing the energy [20].

The high-resolution structure of the PorKN rings presented here reveals that PorK forms a core domain resembling a formylglycine-generating enzyme (FGE). These enzymes catalyse the conversion of a protein cysteine to a protein formylglycine via a proposed copper oxidase mechanism [53]. The two Cys residues in the FGE active site are missing in PorK, indicating that it cannot perform this function. The PorK-specific portion is inserted within the FGE sequence between residue 103 and 261 and is located on the outer side of each subunit forming the outer rim and the interlocking wedge that fits snuggly within the adjacent PorK subunits. Hence one of the functions of the PorK-specific appendage appears to be ring formation. The ring structures in other secretion systems either assemble around a solid core or possess intrinsic properties that facilitate self-assembly. For instance, FliF of Flagella motor self-assembles to form the MS-ring in the inner membrane [54, 55]. Since there is no solid core present in the T9SS, PorK may possess intrinsic properties that enable it to self-assemble and form a ring structure in the periplasm.

MS analysis showed that the *O*-glycosylation sites of PorK and PorN were glycosylated with full length glycans (7 sugars), however in the structure, only the first sugar was resolved at PorK Ser^68^. No sugars were resolved for the two identified glycosylation sites in PorN. The resolution of the structure was not sufficient to resolve the isomeric conformations of the sugars. Presumably, the hidden sugars were not visible due to having high flexibility. The visible sugars participated in multiple hydrogen bonds to different parts of the protein sequence, potentially contributing to the stability of the structure. The role of *O*-glycans on PorK and PorN is unclear, however, SprE, a homolog of PorW in *P. gingivalis*, was also found to be *O*-glycosylated, and this glycosylation was essential for its stability [24]. We therefore predict a similar role for *O*-glycosylation in PorK and PorN.

The structure of PorN was similar to the published crystal structure [56], with the following notable exceptions. The “cog” was displaced by ∼13 Å, and the N-terminal segment (Leu^50^-Asn^67^) as well as the C-terminal segment (Arg^263^-Lys^303^) were resolved here for the first time. PorN binds deeply within a cavity produced by two PorK subunits suggesting that in the ring assembly process, PorK multimerization (at least dimer) may precede the incorporation of PorN subunits. The stability of the PorKN interaction is underscored computationally with a very low estimated K^d^, and experimentally by the ability of the PorKN rings to withstand the harsh purification protocol, including overnight centrifugation in 30% w/v CsCl and short periods in 0.5% DDM, 500 mM NaCl and 1 M urea. Together, these data indicate a solid ring under physiological conditions. The cog-like projections of PorN in the periplasm suggest a mechanism by which the PorLM motor could transfer its rotational energy. AlphaFold modelling of PorM with PorN positioned PorM between the cog-like projections of PorN. Since PorLM motors have been proposed to exhibit rotary movement [20], the rotation of PorM rotors interlocked with PorN cogs would cause the whole PorKN ring to rotate.

PorG co-purifies with the PorKN rings, and we found it to associate with PorK via a disulfide bond between conserved cysteine residues. The inability to resolve PorG in the structure may be due to its low stoichiometry, small size and/or flexibility. Disruption of the disulfide bond made the T9SS non-functional, however the rings now comprising only PorK and PorN appeared similar in structure to the native PorKN rings. This suggests that the disulfide bond with PorG plays a crucial mechanistic role in the function of the T9SS rather than in the assembly of the PorKN rings. PorG has three periplasmic loops. Two of these are the more common short loops of 3-5 aa that are modelled to be buried amongst the hydrophilic phospholipid head-groups, but one loop, Ser^121^-Arg^131^ and the C-terminal extension are long enough to traverse the gap and interact with the surface of PorK. The model predicts PorG to be located on the inside edge of PorK. This location is supported by our previously observed Lys-Lys crosslinks between the periplasmic loop of PorG (Lys^125^) and Lys^480^ and Lys^483^ of PorK[19], with a modelled distance of only 7 Å and 15 Å respectively.

We also demonstrate that the Attachment Complex, consisting of PorU, PorV, PorQ, and PorZ, associates with the PorKN rings. Using immunogold and cryo-ET, we show that this complex is co-located directly above the PorKN rings. Recently, Song *et al* solved a low-resolution *in-situ* structure of the *P. gingivalis* T9SS [21]. An 8-fold symmetry ring of cell surface densities were observed directly above the PorKN periplasmic rings. Based on the observation that these surface densities disappeared in a *sov* mutant, the authors proposed that the surface densities were individual Sov subunits. Interestingly, the densities were also absent in *porL* and *porM* mutants [21]. Our data however, show that Sov is properly located in the OM in *porL* and *porM* mutants maintaining its functional interaction network including PorV, PorW, PorK and PorN (**Extended Data** Fig 7). Similarly, a recent study showed that SprA, the Sov orthologue in *F. johnsoniae*, maintains its complex with PorV and SprE (PorW orthologue) in the *gldL* (*porL)* mutant [24]. Therefore, if the surface densities observed in the *in-situ* structure by Song et al were Sov then those densities shouldn’t disappear in the *porL* mutant. These results suggest that the cell surface rings are unlikely to be the Sov translocons alone. PorU and PorZ of the Attachment Complexes are also T9SS cargo proteins that cannot be secreted to the cell surface in any T9SS component mutants such as *sov*, *porL* and *porM* and therefore it is likely that the observed cell surface densities include the Attachment Complexes, which are of similar overall dimensions to Sov.

Collectively, based on our results we propose the following. PorG OMPs are symmetrically arranged on top of the flat PorK ring surface secured firmly via disulfide bond linkages with the PorK subunits. The OM β-barrels PorQ (bound to PorZ) and PorV (bound to PorU) are also positioned above PorK monomers in close proximity to PorG (**Fig 6C**). The rotation of the PorKN rings would cause the tethered PorGs and the associated Attachment Complexes to also rotate. Recently, it was proposed that energy is needed to detach cargo bound PorV from the Sov translocon [24]. The loaded Sov-PorV-cargo complex is likely to be positioned near the ring via dynamic interaction with the Sov/PorN bridging protein PorW [23] leading to our proposed model of T9SS secretion and attachment in *P. gingivalis* (**Fig 6D**).

In the model the energy of rotation provided by the T9SS PorLM motors is conveyed to the PorKN rings. The bent PorM rotors fit into the PorN cogs to drive the rotation of the ring along with the disulfide-bonded PorG and associated Attachment Complexes. With Sov in close proximity to the rotating ring it positions the PorV-cargo in the path of the rotating PorG. When the PorG and Attachment Complexes are driven past Sov-PorV, PorG dislodges PorV, pushing it away from Sov, and simultaneously pulling the cargo through Sov’s lateral gate thereby completing secretion (**Fig 6D**). The extracted PorV-cargo complex may then bind to the Attachment Complex also rotating past leading to PorU-catalysed CTD cleavage and A-LPS attachment of the cargo. This represents an efficient and coordinated model of secretion and attachment, since the cargo substrates would be transferred directly from Sov to the Attachment Complexes by the action of PorG, rather than utilizing the less efficient shuttle model that we previously suggested [8, 25]. Our previous data can nevertheless be explained whereby in the absence of PorU or PorZ of the Attachment Complexes, the PorV cargo complexes accumulate [25]. In these mutants, PorG can presumably still function to extract the PorV-cargo complexes from Sov but without functional Attachment Complexes the PorV-cargo cannot be processed and therefore accumulate. The model also allows for one or more Sov translocons per ring.

In conclusion, we present a high-resolution structure of the T9SS PorKN rings and demonstrate a close association with other components of the T9SS, in particular PorG and the Attachment Complexes. Together, these findings provide novel insights into the molecular mechanism of T9SS secretion and attachment in *P. gingivalis*.

## Funding

This work was supported by the Australian Government, Department of Industry, Innovation and Science (Grant ID: 20080108), the Australian National Health and Medical Research Council (Grant IDs: 1193647, 1123866 and APP1196924), the Australian Research Council (Grant ID: DP200100914) and the Australian Dental Research Foundation (Grant ID: 492-2019) and a Cumming Global Centre for Pandemic Therapeutics, Foundation Grant.

## Author contributions

D.G.G., conceptualisation, methodology, data acquisition, investigation, formal analysis, data curation, funding acquisition, and writing—original draft; E.H., data acquisition, investigation, formal analysis, data curation, C.M, M.M, S.V., D.G., data acquisition and writing—review and editing, C.S., L.Z, methodology and writing—review and editing; P.D.V., conceptualisation, investigation, data curation, supervision, funding acquisition and writing—review and editing; E.C.R., conceptualisation, supervision, funding acquisition and writing—review and editing. All authors have read and agreed to the published version of the manuscript.

## Acknowledgments

We acknowledge the use of the Mass Spectrometry and Proteomics Facility and the Ian Holmes Imaging Centre both at the Bio21 Institute, The University of Melbourne, Australia

---

**Extended Data Fig 1:**
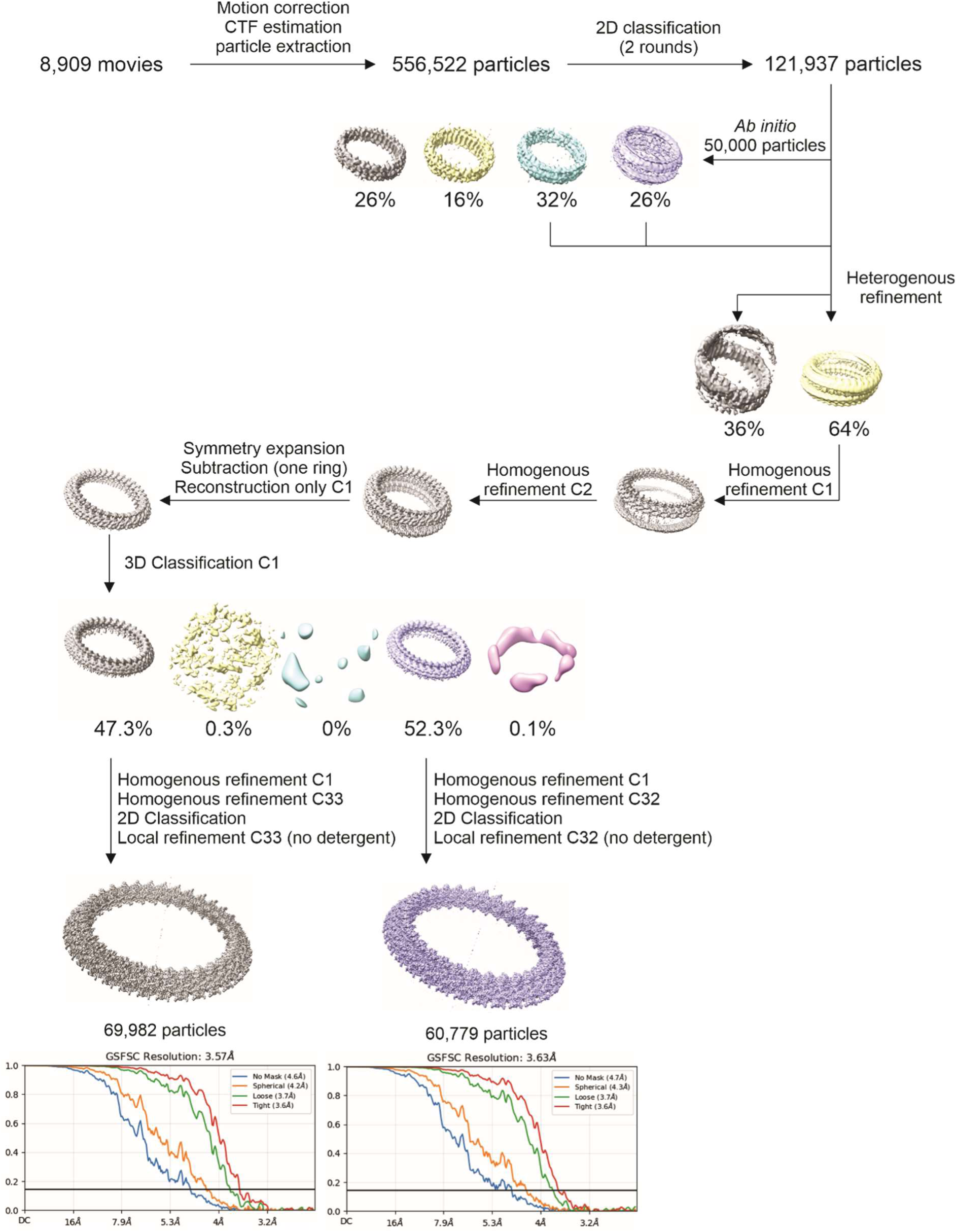
Workflow for single particle analysis of PorKN rings.

**Extended Data Fig 2:**
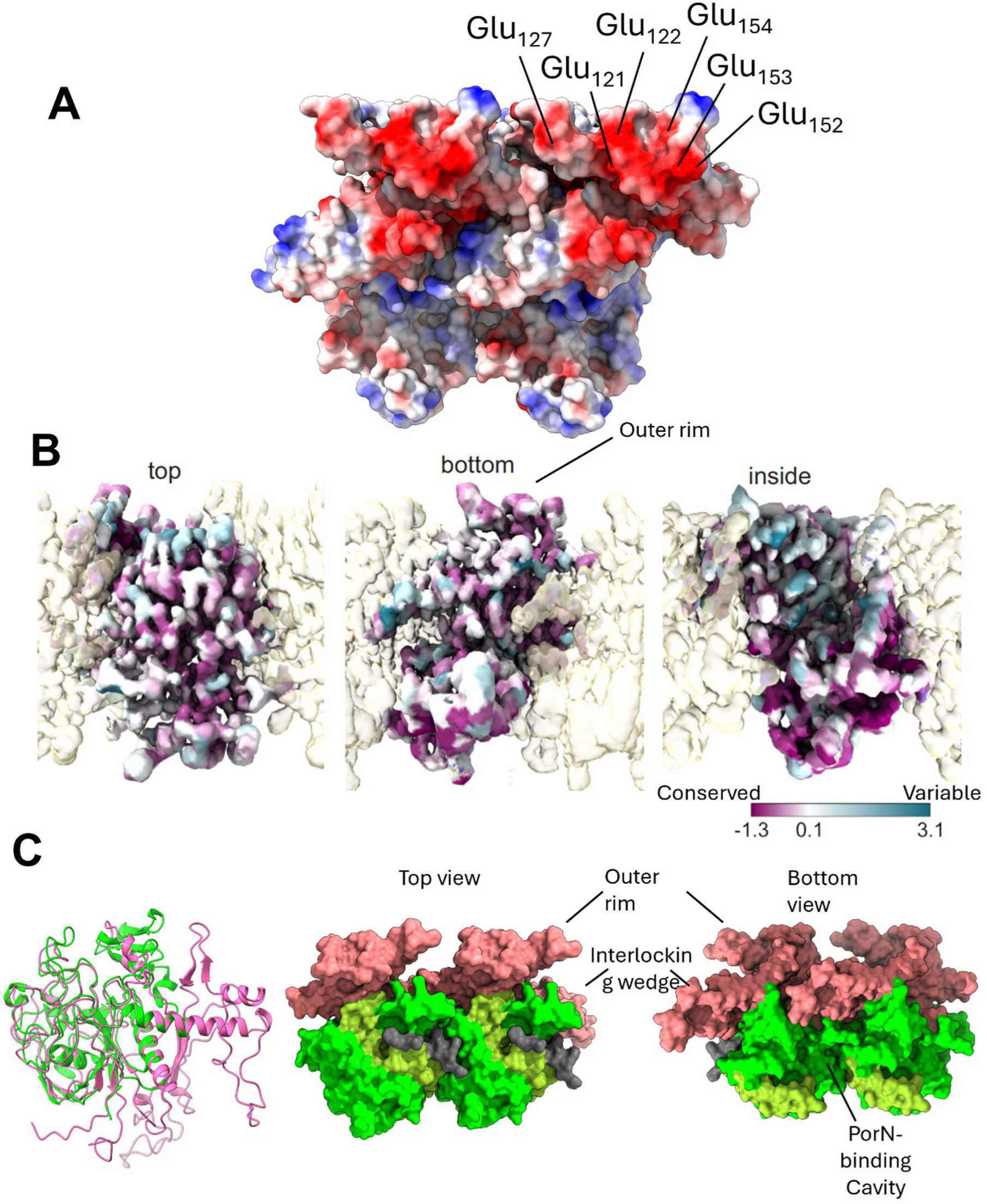
Analysis of PorK and PorN proteins. (A) Surface charge of PorK_2_N_2_ unit (red = negative, blue = positive, white = neutral). The negatively charged residues of the three-toed claw of the outer rim are shown. (B) Conservation map of the PorKN heterodimer showing highly conserved outer rim and the PorN cog. (C) Left-Overlay of PorK (pink) and FGE sulfatase (green). Location of the FGE domain within a PorK_2_ unit. The FGE domains span residues 50-103 (light green) and 261-471 (green), while the PorK-specific domain spanned residues 104-260 (salmon).

**Extended Data Fig 3:**
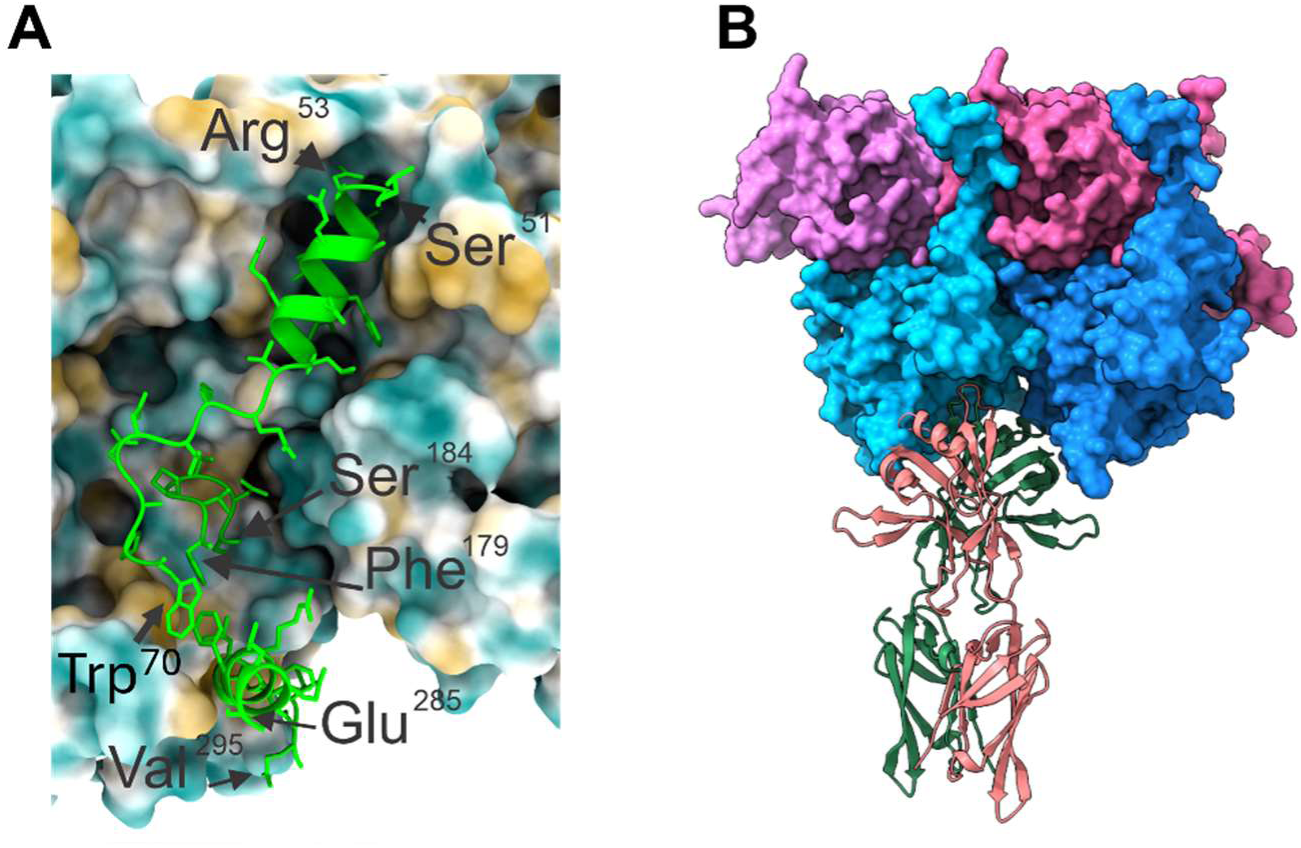
Analysis of PorK-PorN and PorN-PorM interaction. (A) A zoomed-in view of the PorN interaction with a pair of PorK subunits shows three distinct interface regions: Leu^50^-Trp^70^, Phe^179^-Ser^184^, Glu^285^-Val^295^. (B) AlphaFold3 prediction of PorM_2_ C-terminal domains interacting with PorK_2_PorN_2_. PorK domains are in shades of pink, PorN domains in shades of blue while the two PorM subunits are shown in green and salmon.

**Extended Data Fig 4:**
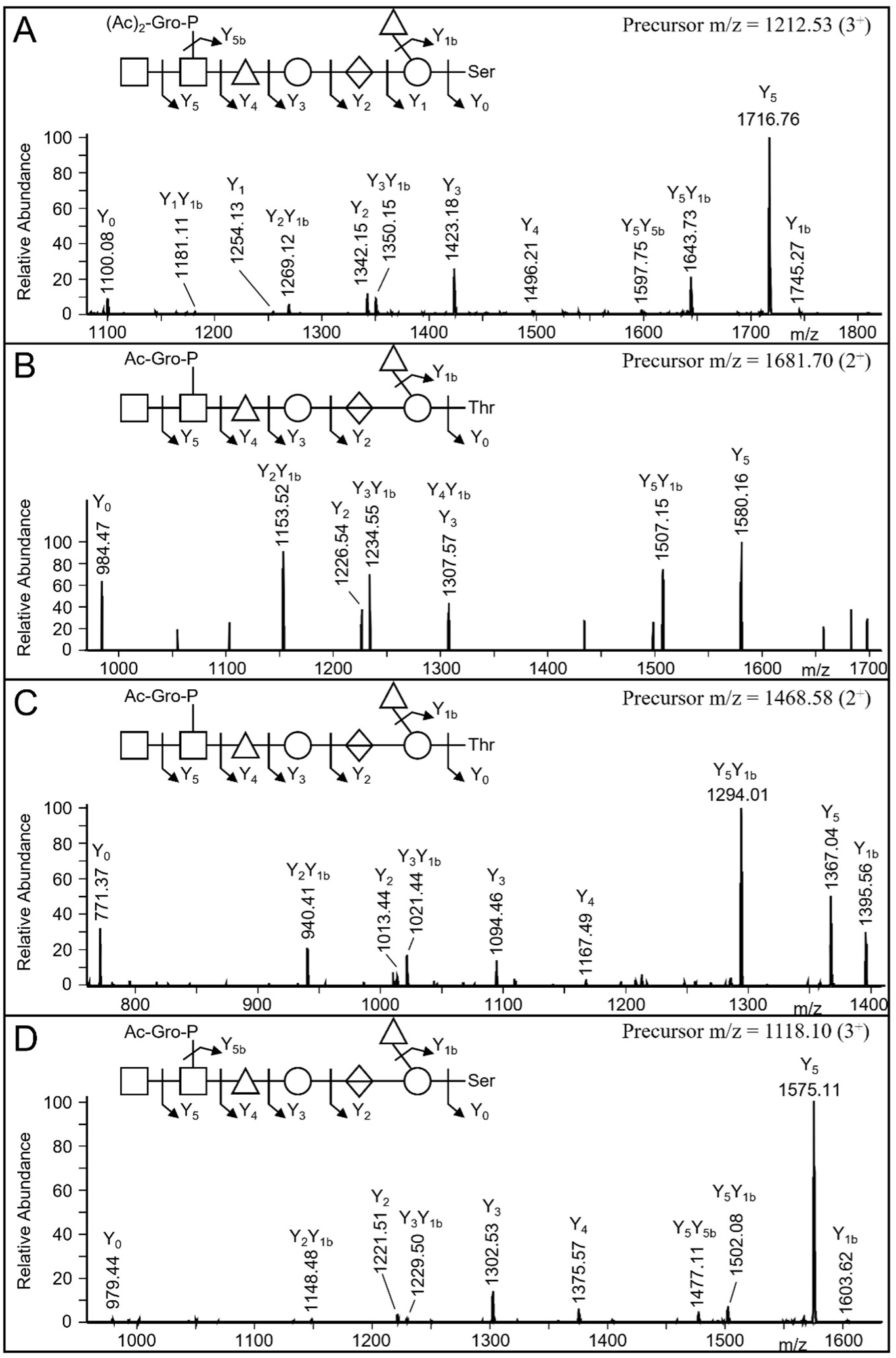
CID spectra of tryptic peptides modified with *O-*glycans. (A) PorK peptide GSIVLGNKEAD<u>S</u>LWGIPAESR modified with the Δmass 1436 Da *O-* glycan. (B) PorK peptide D<u>T</u>AFVDESGNIISETITR modified with the Δmass 1394 Da *O-* glycan. (C) PorN peptide DGLFVTQD<u>T</u>TWMK modified with the Δmass 1394 Da *O-*glycan. (D) PorN peptide ALAEYCPTPD<u>S</u>MKMESK modified with the Δmass 1394 Da *O-*glycan. The CID spectra were triggered by the presence of HexNAc (m/z 204.09) in the corresponding HCD spectra. For all spectra, the labelled ions are 2^+^. The structure of the glycans are shown above each spectrum together with the fragmentation scheme matching the observed fragment ions. Sugar symbols follow the SNFG as described in Materials and Methods.

**Extended Data Fig 5:**
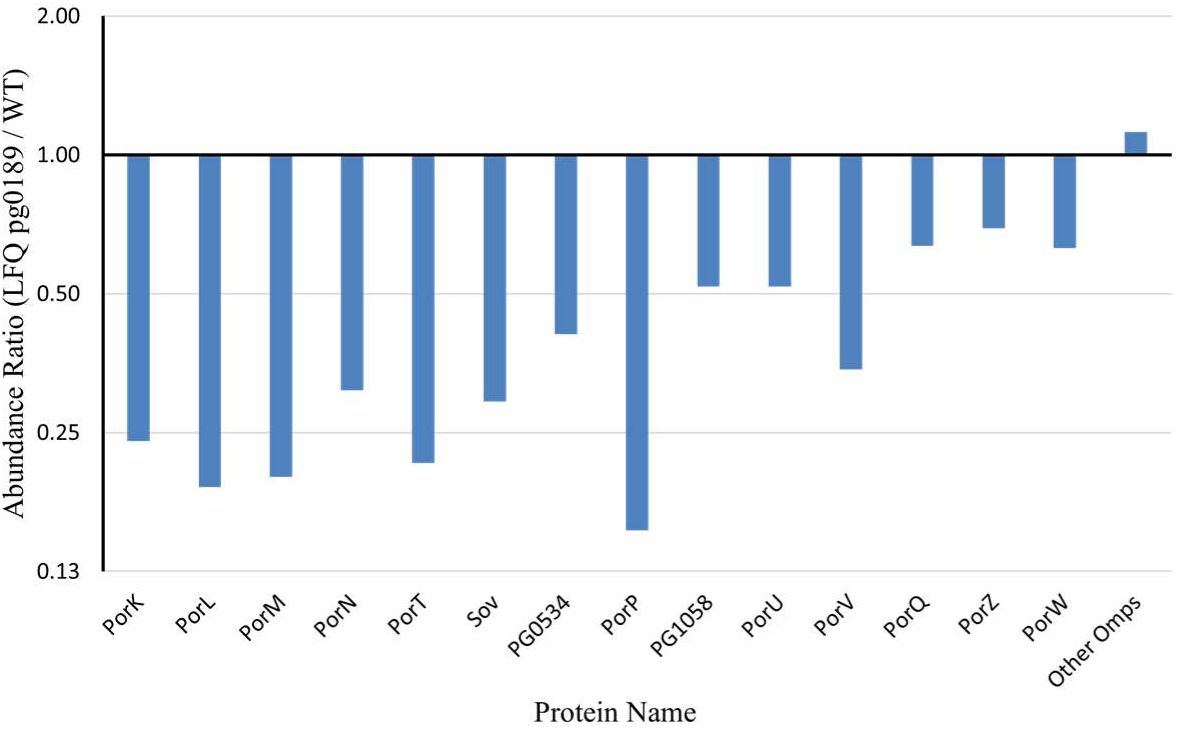
Abundance of the T9SS proteins in the *porG* mutant relative to WT. Cell lysates from WT and *porG* mutant were electrophoresed on an SDS-PAGE for a short period. The band consisting of all the proteins was excised and subjected to mass spectrometry analysis. The proteins were quantified using MaxQuant software.

**Extended Data Fig 6:**
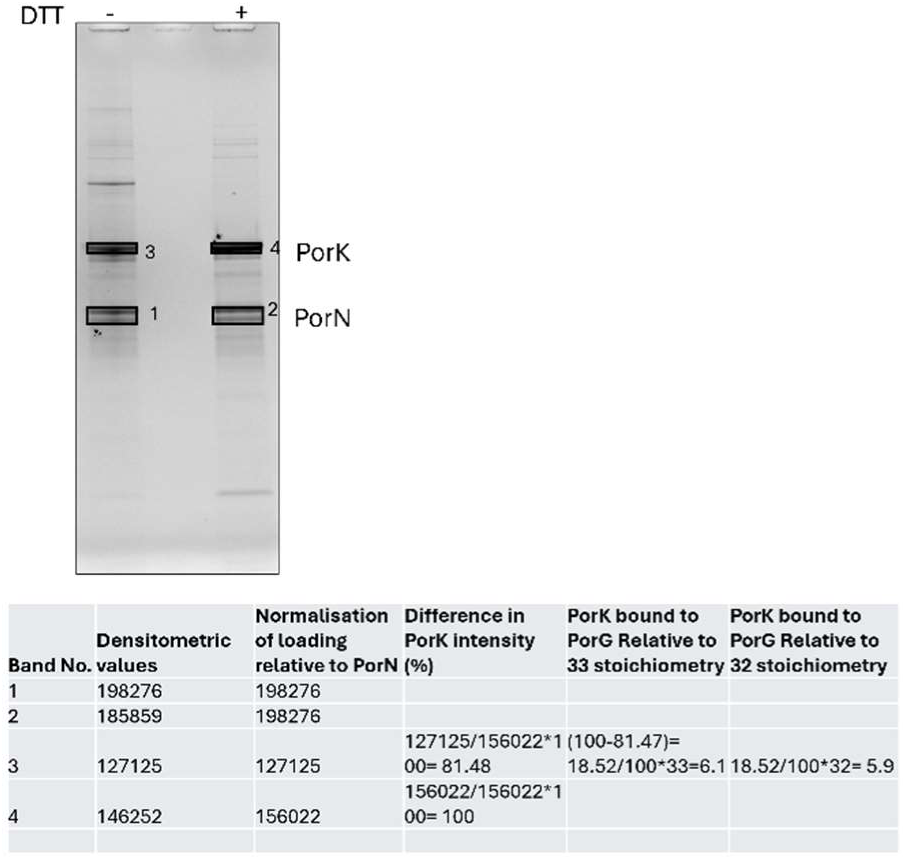
Stoichiometry of PorG. SDS-PAGE gel of PorKNG complex electrophoresed under reducing and non-reducing conditions. Bands labelled 1, 2, 3 and 4 were quantified by imageJ software (n=2).

**Extended Data Fig 7:**
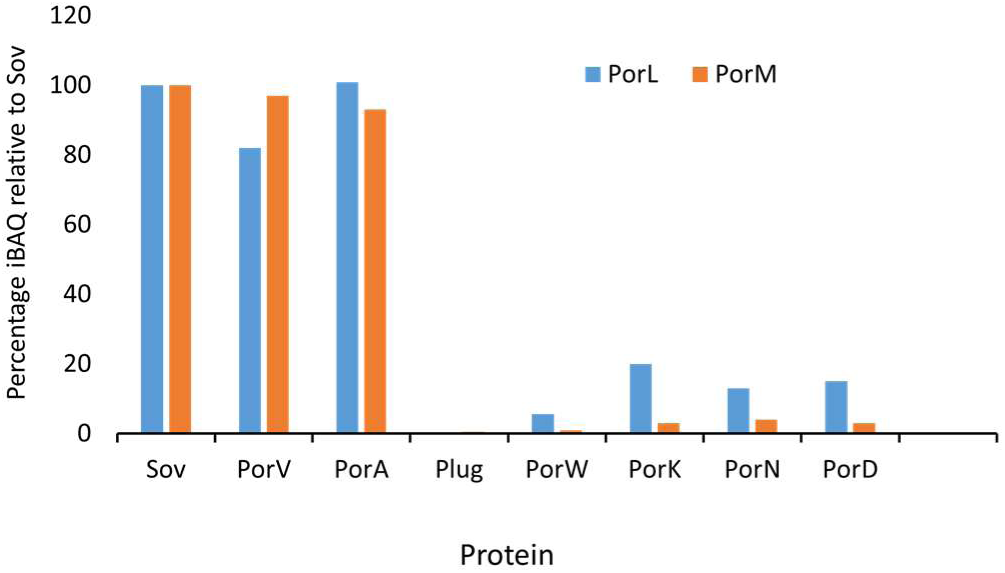
Sov interactome in *porL* and *porM* mutants. The *P. gingivalis porL* and *porM* mutants were subjected to coimmunoprecipitation using Sov-specific antibodies. The *sov* mutant was used as a negative control. The immunoprecipitated samples were digested with trypsin, analyzed by mass spectrometry, and quantified using MaxQuant software. Percentage of abundance relative to Sov using iBAQ intensities was plotted.

## Notes

### Competing Interest Statement

The authors have declared no competing interest.

