## Supplementary Table 1 for "Near-atomic structure of the PorKN rings, disulfide bonded to PorG and bound to Attachment Complexes, provide mechanistic insights into the type IX secretion system"

| **Table S1. Cryo-EM statistics** |  |  |
| --- | --- | --- |
|  | **porKN ring C33** | **porKN ring C32** |
|  | **PDB 9MYJ/ EMD-48741** | **EMD-48722** |
| **Data collection** |  |  |
| Microscope | FEI Titan Krios | FEI Titan Krios |
| Voltage (kV) | 300 | 300 |
| Camera | Gatan K3 infinium | Gatan K3 infinium |
| Mode | Counting | Counting |
| Magnification | 64,000 | 64,000 |
| Pixel size (Å) | 1.32 | 1.32 |
| Electron dose (e–/Å^2^) | 54 | 54 |
| Defocus range (μm) | -0.6 to -2.0 | -0.6 to -2.0 |
| Exposure (s) | 5.3 | 5.3 |
| **Reconstruction** |  |  |
| Software | CryoSPARC-4.1.1 | CryoSPARC-4.1.1 |
| Frames | 40 | 40 |
| Symmetry | C33 | C33 |
| Initial particle images (no.) | 556,522 | 556,522 |
| Final particle images (no.) | 69,982 | 60,779 |
| Map resolution, FSC_0.143_ (Å) | 3.57 | 3.63 |
| Map-sharpening B factor (Å) | -110.1 | -108.0 |
| **Refinement** |  |  |
| Software | Phenix-1.20.1 |  |
| **Model composition** |  |  |
| Non-hydrogen atoms | 5899 |  |
| Protein residues | 700 |  |
| **R.m.s. deviations** |  |  |
| Bond lengths (Å) | 0.004 |  |
| Bond angles (°) | 0.717 |  |
| **Validation** |  |  |
| MolProbity score | 2.29 |  |
| Clashscore | 11.97 |  |
| Poor rotamers (%) | 1.94 |  |
| **Ramachandran plot** |  |  |
| Favored (%) | 92.1 |  |
| Allowed (%) | 7.76 |  |
| Disallowed (%) | 0.14 |  |
